# OncoSpot enables benchmarking of cancer spatial transcriptomics methods across tumor evolutionary modes

**DOI:** 10.64898/2026.11.01.763496

**Authors:** Pankaj Bhattarai, Weiman Yuan, Xin Maizie Zhou, Xian Mallory

## Abstract

Spatial transcriptomics (ST) is increasingly used to map clonal architecture within tumors, yet methods for analyzing cancer ST data are difficult to evaluate because real tumors lack ground truth for copy-number alterations (CNAs), subclonal identity, and malignant cell fraction. Existing ST simulators model normal tissue and do not represent tumor evolution, CNA-driven expression, or the spatial organization of malignant subclones. We present OncoSpot, a cancer-aware simulator of Visium-like spot-based ST data. OncoSpot combines a spatial evolutionary scaffold from DEMON, CNAs generated along the resulting clonal phylogeny with CNAsim, cancer-type-specific malignant transcriptional programs, and spatially clustered nonmalignant cell types, and aggregates cells into spots with known malignant fraction, subclone, and copy-number profiles. We demonstrate OncoSpot on high-grade serous ovarian carcinoma, colorectal cancer, and hepatocellular carcinoma, which represent three distinct modes of tumor evolution. Simulated subclones were spatially clustered and showed stronger phylogeny–space concordance than those generated by scCube under matched subclone populations. Benchmarking CopyKAT, CancerFinder, Clonalscope, and RCTD revealed cancer-dependent performance: Clonalscope recovered copy-number signals more accurately in ovarian and colorectal cancer, whereas CopyKAT performed better in hepatocellular carcinoma, reflecting both the assumptions of each method and the copy-number landscapes of the three cancers. OncoSpot’s ground truth also exposed a failure mode in Clonalscope’s malignant spot identification, in which tumor and normal labels were inverted, and allowed us to trace its cause. These method-dependent behaviors would be difficult to detect in real data. OncoSpot provides a framework for benchmarking CNA inference, subclone detection, malignant spot identification, and normal spot deconvolution, and its modular design and guidelines allow extension to additional cancer types. OncoSpot is freely available at https://github.com/compbio-mallory/OncoSpot.

## 1 Introduction

Spatial transcriptomics (ST) enables the measurement of gene expression while preserving its location within tissue, providing a framework for investigating molecular heterogeneity in its spatial context [1]. This capability is particularly important in cancer, where tumor evolution produces genetically distinct malignant subclones that coexist with diverse nonmalignant populations within a spatially organized microenvironment. The spatial distribution of malignant subclones reflects their evolutionary histories, local expansion, and interactions with surrounding tissue. Spatial genomic studies have revealed complex patterns of clonal coexistence and segregation, as well as associations between genetic alterations, transcriptional phenotypes, and local microenvironmental composition [2–4]. Copy-number alterations (CNAs), which involve gains or losses of genomic regions, constitute an important source of intratumoral genetic heterogeneity and can influence gene expression through altered genomic dosage. Spatially resolved transcriptomics therefore offers an opportunity to investigate the relationship between clonal evolution, genomic alterations, and transcriptional heterogeneity within intact tumor tissue [5, 6].

However, reconstructing malignant genomic states from spatial transcriptomic measurements remains challenging. RNA abundance does not directly measure DNA copy number, and observed expression reflects multiple biological processes, including genomic dosage, cellular identity, transcriptional programs, and microenvironmental influences. Pan-cancer single-cell studies have identified recurrent malignant transcriptional programs, including proliferation, stress responses, metabolic activity, and epithelial–mesenchymal transition, that can vary independently of clonal genotype [7, 8]. In addition, spot-based technologies such as 10x Genomics Visium capture transcripts from multiple cells within each spatial location. Consequently, observed expression represents a mixture of malignant and nonmalignant populations, potentially including several malignant subclones. Differences in tumor purity, cell-type composition, transcriptional activity, and sequencing depth can therefore obscure CNA-associated signals or produce expression patterns that resemble genomic changes. Distinguishing these sources of variation is essential for accurately recovering malignant regions, clonal identities, and copy-number profiles from spatial expression data [3, 9, 6].

Several computational approaches have been developed to infer malignant genomic structure from transcriptomic data. STARCH uses spatial expression measurements to infer copy-number profiles and tumor clones, whereas CalicoST integrates transcript counts, allele-specific information, and spatial relationships to reconstruct CNAs and tumor phylogeography [5, 6]. Clonalscope provides another approach for detecting malignant subclones from copy-number-associated signals in single-cell and spatial omics data [10]. Related methods address malignant-cell identification, including CopyKAT, CancerFinder, and scMalignantFinder [11–13]. DeepMalignant further combines cancer-associated expression features with CNA-informed cellular relationships through graph representation learning and has been applied to spatial breast cancer samples with histological comparison [14]. Beyond malignant classification and CNA inference, reference-based deconvolution methods such as RCTD and cell2location estimate the cellular composition of spatial measurements, addressing another important consequence of mixed-cell spot capture [15, 16]. These approaches address complementary analytical tasks, but their predictions depend on different modeling assumptions, input requirements, and properties of the underlying tumor samples.

A fundamental challenge in evaluating these methods is the limited availability of spatially resolved ground truth. Histopathological annotations can identify tumor-enriched regions, while matched DNA-sequencing or spatial genomic assays provide more direct evidence of genomic alterations. For example, spatial DNA-sequencing and integrated genomic mapping approaches have demonstrated the feasibility of resolving distinct cancer clones and their relationships to tissue architecture [3, 2]. Nevertheless, obtaining matched cell identities, clonal ancestry, copy-number states, and transcriptomic measurements at the same spatial locations remains technically demanding. Furthermore, different computational methods may be affected differently by tumor purity, clonal diversity, CNA burden, and spatial mixing. A recent systematic benchmark of nine CNA inference methods across 69 spatial transcriptomics sections from six cancer types found that no single method consistently outperformed the others across all evaluated tasks [17]. These findings highlight the importance of evaluating computational methods under well-defined biological and technical conditions rather than relying exclusively on aggregate performance across heterogeneous real datasets.

Simulation provides a complementary strategy by generating data with explicit ground truth and allowing individual properties of the data-generating process to be controlled. Spatial transcriptomics simulators have made substantial progress in reproducing expression distributions and spatial patterns. SRTsim supports reference-based and reference-free simulations designed to preserve spatial expression characteristics, while scCube generates spatially structured cell or spot populations and supports benchmarking of deconvolution, imputation, and spatial reconstruction [18, 19]. A systematic evaluation of single-cell and spatial transcriptomics simulators has further demonstrated that simulation accuracy, functionality, and scalability vary across methods and experimental settings [20]. These tools provide valuable resources for assessing expression-based analyses. However, evaluating cancer genomics methods additionally requires a representation of the relationships among malignant subclone ancestry, inherited genomic alterations, spatial cell organization, and the resulting transcriptomic observations.

Recent cancer-focused simulation frameworks have begun to address these requirements from complementary perspectives. STGBench generates spatially indexed DNA-sequencing outputs and matched transcriptomic data from specified genomic alteration patterns, supporting benchmarking of spatial CNA and mutation analysis [21]. stCNASim introduces allele-aware spatial RNA-sequencing simulation and examines CNA inference under varying technical conditions and spatial architectures [22]. These developments emphasize the growing importance of genomic ground truth in spatial transcriptomics benchmarking. A complementary direction is to connect spatially structured tumor evolution and inherited clone-specific CNAs with cancer-dependent transcriptional states, nonmalignant cell populations, and spot-level observations within a unified cell-to-spot simulation framework. Such a framework can provide the known cellular and genomic annotations needed to investigate multiple downstream tasks using the same underlying simulated tissue.

Here, we introduce OncoSpot, a cancer-oriented simulator that integrates deme-based tumor evolution, malignant subclone phylogeny, clone-specific CNA profiles, reference-calibrated gene expression, and Visium-like spatial observation. OncoSpot uses DEMON to generate spatially structured evolutionary histories and a driver phylogeny, which is subsequently supplied to CNAsim to simulate inherited copy-number alterations along the retained malignant subclone tree [23, 24]. The resulting spatial and genomic information is combined with cancer-specific malignant transcriptional programs and nonmalignant cell-type distributions to generate cell-level RNA counts. These counts are then aggregated into Visium-like spots with empirical sequencing-depth characteristics. Importantly, OncoSpot retains the underlying cellular identities, spatial coordinates, malignant subclone assignments, cell-type compositions, and gene-level copy-number profiles, providing matched ground truth at both cell and spot resolution. This design enables direct evaluation of downstream predictions against the known generating states without requiring the simulated expression data to uniquely determine those states.

We demonstrate OncoSpot using three cancer-specific simulation configurations representing high-grade serous ovarian carcinoma (HGSOC), colorectal cancer (CRC), and hepatocellular carcinoma (HCC), representing three of the four evolutionary modes modeled by DEMON [23]: branching evolution (HGSOC; invasive glandular growth), progressive diversification (CRC; gland fission), and effectively almost neutral evolution (HCC; boundary growth). These configurations incorporate distinct spatial growth scenarios and cancer-specific transcriptomic references rather than assuming a universal tumor architecture. We evaluate spatial autocorrelation of malignant subclones and the correspondence between phylogenetic and spatial relationships, including comparisons with scCube under matched malignant subclone populations. We further benchmark malignant spot identification using CopyKAT, CancerFinder, and Clonalscope; malignant subclone recovery and copy-number inference using Clonalscope and CopyKAT; and nonmalignant cell-type deconvolution using RCTD. Across the three cancer configurations, OncoSpot generates spatially clustered malignant subclones with positive phylogeny–space concordance, while downstream inference performance varies substantially across cancer types and analytical tasks. These analyses illustrate the utility of OncoSpot, whose explicit cellular and genomic ground truth reveals method-dependent behavior under controlled conditions. Beyond the three cancer types evaluated here, OncoSpot provides detailed guidelines for users to configure new cancer types, enabling simulation and benchmarking across a broader range of tumors.

## Results

### Method overview

OncoSpot integrates spatial tumor evolution, copy-number alteration (CNA) simulation, and reference-based gene expression modeling to generate spatial transcriptomics data with known cellular and clonal ground truth (**Fig. 1**). First, DEMON generates a spatially structured deme population and its associated driver phylogeny, which is subsequently supplied to CNAsim to simulate clone-specific CNA profiles along the evolutionary tree (**Fig. 1A**). The DEMON-derived spatial scaffold and CNAsim-generated CNA profiles are then integrated with cancer-specific transcriptional programs and reference-based expression models to simulate cell-level gene expression while preserving spatial and clonal relationships (**Fig. 1B**). The resulting tissue contains spatially organized malignant subclones and nonmalignant cells, each with known spatial coordinates, cellular identity, and gene expression profiles (**Fig. 1C**). These cell-level expression profiles are aggregated into Visium-like spatial spots to generate spot-level gene-count matrices while retaining the underlying cellular composition and copy-number ground truth (**Fig. 1D**). Finally, the generated datasets enable benchmarking of downstream computational methods for malignant spot identification, recovery of malignant clonality, total copy-number inference, and cell-type deconvolution (**Fig. 1E**).

**Figure 1:**
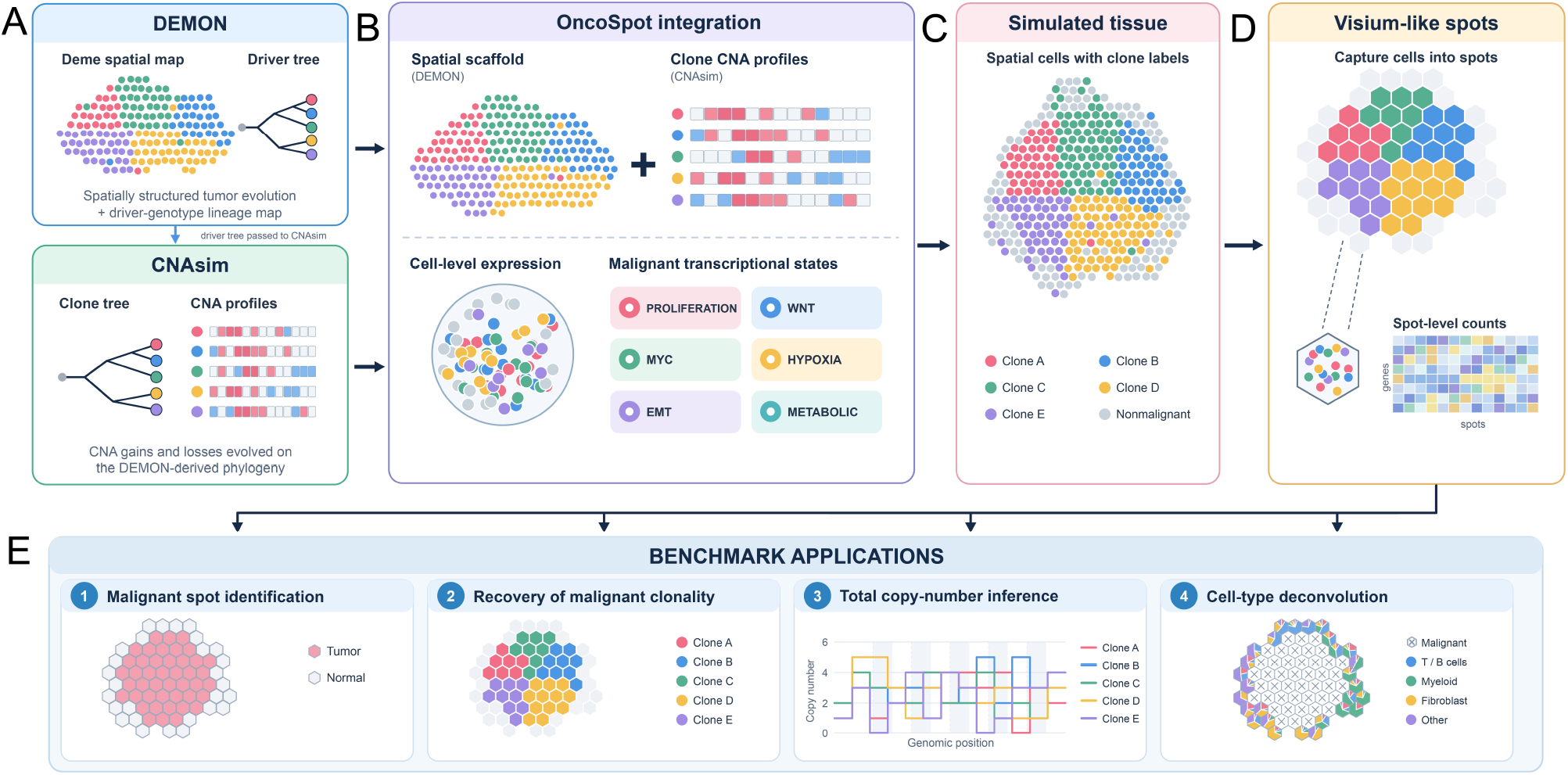
Overview of the OncoSpot simulation framework. **(A)** DEMON simulates spatially structured tumor evolution, generating a deme-based spatial map and driver phylogeny. The resulting phylogeny is supplied to CNAsim, which simulates clone-specific copy-number alterations along the evolutionary tree. **(B)** OncoSpot integrates the DEMON-derived spatial scaffold with CNAsim-generated CNA profiles and cancer-specific transcriptional programs to simulate cell-level gene expression. **(C)** The resulting simulated tissue contains spatially organized malignant subclones and nonmalignant cells, with known cell identities, spatial coordinates, CNA profiles, and gene expression. **(D)** Individual cells are captured and aggregated into Visium-like spatial spots, producing spot-level gene-count matrices while preserving cellular composition and clonal ground truth. **(E)** The simulated datasets support benchmarking of malignant spot identification, malignant clonality recovery, total copy-number inference, and cell-type deconvolution.

### Simulated spatial transcriptomics datasets across three cancer types

We evaluated OncoSpot-generated spatial transcriptomics datasets representing high-grade serous ovarian carcinoma (HGSOC), colorectal cancer (CRC), and hepatocellular carcinoma (HCC). Each simulation comprised 15,000 cells, from which 418, 409, and 414 Visium-like spots were obtained for HGSOC, CRC, and HCC, respectively. **Supplemental Section S1** listed details of these tumors’ evolution architectures and their biological interpretations.

The simulated datasets retained the exact cellular composition of each spot, including malignant and nonmalignant cell identities, malignant subclone assignments, and gene-level absolute total copy numbers. The observed spots contained 2,107, 2,058, and 2,065 malignant cells in HGSOC, CRC, and HCC, respectively, representing 10, 11, and 10 malignant subclones.

Using a malignant-cell fraction of at least 50% to define tumor-dominated spots, the datasets contained 219 HGSOC, 208 CRC, and 222 HCC tumor spots. These known cellular and genomic annotations enabled direct evaluation of malignant spot identification, subclone recovery, and copy-number reconstruction.

### Spatial autocorrelation and comparison with scCube

We evaluated whether OncoSpot generates spatially clustered malignant subclones using join-count statistics and per-clone Moran’s *I* [25] (Eqs. (25) and (28)). The analysis included 7,500 malignant cells per cancer, representing five retained subclones in HGSOC, six in CRC, and five in HCC.

All three cancer datasets exhibited substantial enrichment of neighboring malignant cells belonging to the same subclone compared with random clone-label placement (**Table 1**). In HGSOC, 84.1% of neighboring malignant-cell pairs belonged to the same subclone, compared with 44.1% expected under random placement, corresponding to a 1.91-fold enrichment. CRC exhibited 67.5% same-clone neighboring pairs compared with a random expectation of 37.8%, representing a 1.79-fold enrichment. HCC showed the strongest local clustering among the three simulated datasets, with 88.0% same-clone neighboring pairs compared with 35.1% expected under random placement, corresponding to a 2.50-fold enrichment. Permutation testing confirmed significant spatial enrichment in all three datasets (*P* = 0.001, the smallest attainable value with 999 permutations).

**Table 1:**
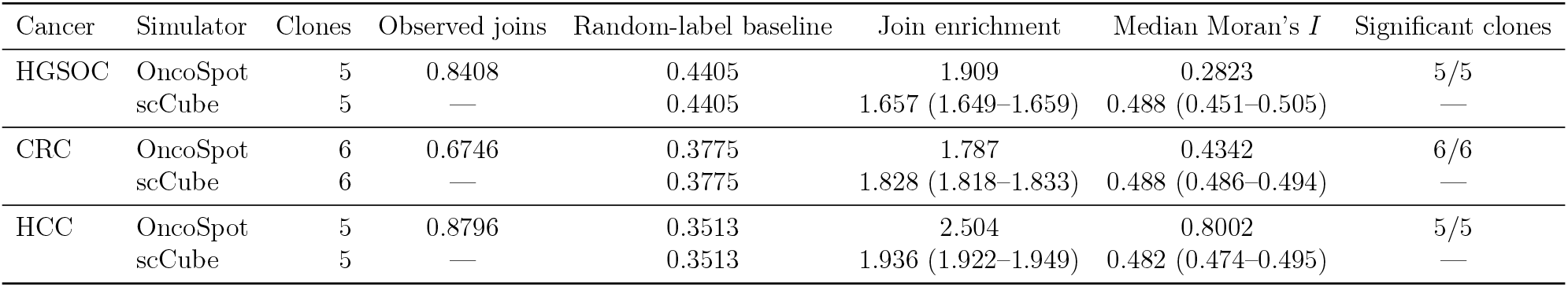
Spatial autocorrelation of malignant subclones and comparison with scCube. OncoSpot values are from one finalized realization per cancer, and scCube join enrichment and median clone-specific Moran’s *I* are medians across ten reference-free spatial realizations (interquartile ranges in parentheses). Each simulator uses 7,500 malignant cells per cancer with matched clone identities and abundances. Observed joins denote the fraction of undirected edges connecting same-clone cells on a symmetrized eight-nearest-neighbor graph; the random-label baseline is calculated from clone abundances (Eq. (26)). Significant clones have positive binary-indicator Moran’s *I* and Benjamini–Hochberg-adjusted one-sided permutation *q <* 0.05 (999 permutations). Dashes indicate quantities not tabulated for individual scCube realizations in the original comparison.

Consistent with these findings, per-clone Moran’s *I* values were positive for every retained malignant subclone. Median Moran’s *I* values were 0.282, 0.434, and 0.800 for HGSOC, CRC, and HCC, respectively. Across the 16 retained malignant subclones, Moran’s *I* ranged from 0.210 to 0.918, and all subclones exhibited significant positive spatial autocorrelation after Benjamini–Hochberg correction (adjusted *q* = 0.001). A subclone was counted as significant when its observed binary-indicator Moran’s *I* exceeded the random-label distribution in a one-sided permutation test and its Benjamini–Hochberg-adjusted *q*-value was below 0.05. Each test used 999 permutations of the clone labels over the fixed spatial graph; the number of cells assigned to each clone was kept constant.

These results demonstrate that OncoSpot generates nonrandom spatial organization of malignant subclones across distinct cancer-specific simulation configurations. We further compared this organization with the reference-free spatial-pattern generator of scCube [19], using identical malignant subclone identities and abundances and ten scCube spatial realizations per cancer. Both simulators generated spatially clustered subclones (**Fig. 2**); all 30 scCube realizations showed significant same-clone join enrichment (permutation *P* = 0.001 for each). The strength of autocorrelation nevertheless varied by cancer (**Table 1**). OncoSpot showed greater same-clone join enrichment in HGSOC (1.909 versus 1.657) and HCC (2.504 versus 1.936), whereas scCube was slightly higher in CRC (1.828 versus 1.787). OncoSpot had the greater median per-clone Moran’s *I* in HCC (0.800 versus 0.482), while scCube had higher values in HGSOC and CRC. Thus, the two simulators exhibit distinct spatial organization under matched clone populations; the differences in these spatial autocorrelation metrics alone do not establish biological realism.

**Figure 2:**
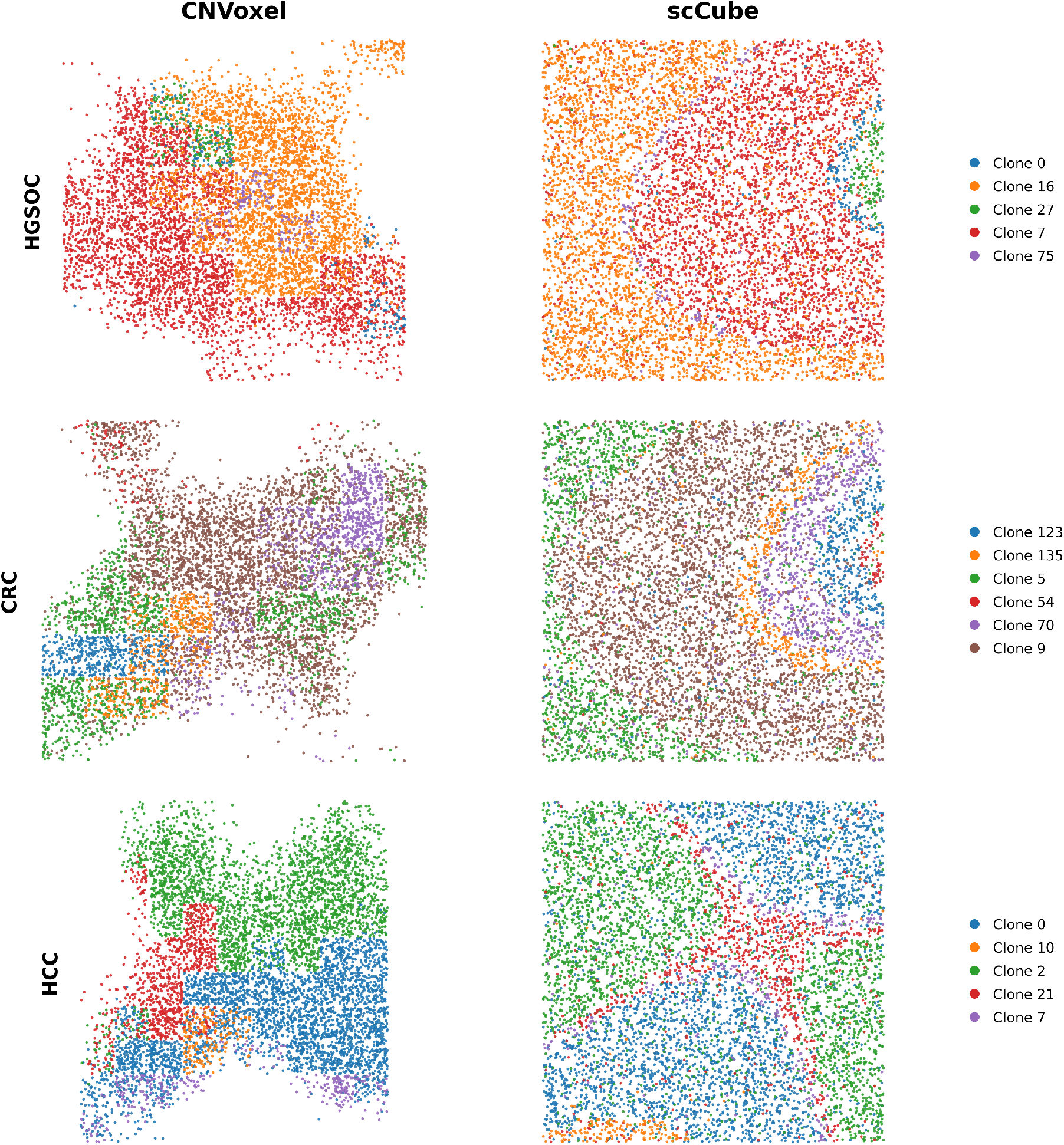
Comparison of malignant subclone spatial organization between OncoSpot and scCube. Spatial distributions of 7,500 malignant cells across high-grade serous ovarian carcinoma (HGSOC), colorectal cancer (CRC), and hepatocellular carcinoma (HCC). Both simulators use identical malignant subclone identities and abundances, with matching colors indicating corresponding subclones within each cancer type. OncoSpot shows the simulated tissue-associated spatial organization, whereas scCube shows a representative realization generated using its reference-free spatial-pattern model. Spatial coordinates are displayed in each simulator’s native coordinate system. Quantitative comparisons of spatial autocorrelation are provided in **Table 1**.

### Phylogeny–space concordance across independent simulations

We next investigated whether the spatial organization of OncoSpot-generated malignant subclones was consistent with their evolutionary relationships. For each of HGSOC, CRC, and HCC, we generated 50 independent simulations and quantified the phylogenetic distance between different dominant malignant subclones occupying adjacent Visium spots. The observed boundary statistic was compared with a null distribution obtained by shuffling malignant subclone identities on the corresponding evolutionary tree, while preserving the spatial clone arrangement and tree topology (Eqs. (29) and (30)).

Of the 50 attempted realizations per cancer, valid standardized concordance scores were obtained for 42 HGSOC, 37 CRC, and 45 HCC datasets. The remaining realizations were excluded because of incomplete simulation outputs, insufficient retained subclone diversity, absence of qualifying cross-subclone boundaries, or zero variance in the permutation null distribution. No realizations were excluded based on the direction or magnitude of their concordance scores.

All three cancer types exhibited predominantly positive concordance scores (**Table 2**). HGSOC had a median *z*-score of 1.149 (interquartile range, 0.712–1.611), with positive scores in 37 of 42 evaluable realizations. CRC had a median *z*-score of 1.413 (interquartile range, 1.000–1.706), with 35 of 37 scores positive. HCC exhibited the largest median concordance score, 1.728 (interquartile range, 1.081–2.007), and all 45 evaluable realizations had positive scores.

**Table 2:**
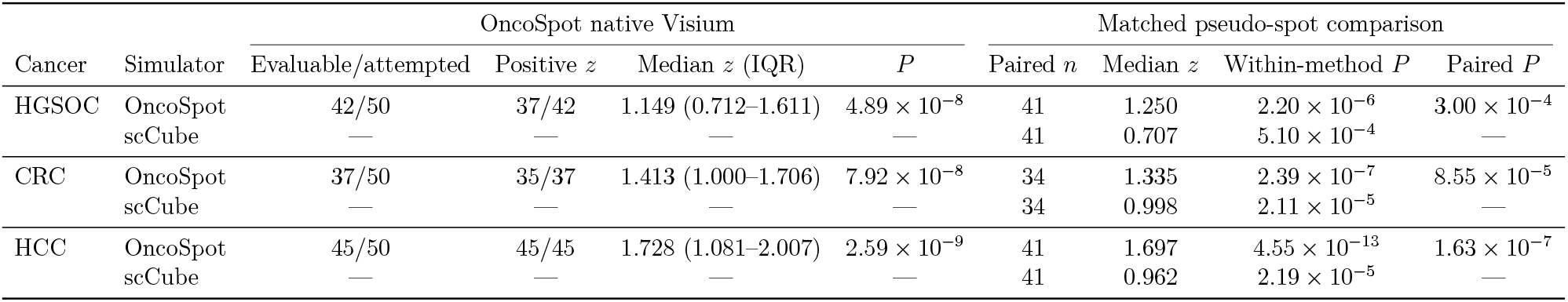
Phylogeny–space concordance in OncoSpot and scCube. The native-Visium analysis evaluates OncoSpot across 50 independent simulations per cancer. The matched comparison uses identical malignant subclone populations and phylogenies with a common malignant-cell-derived pseudo-spot procedure. Within-method *P* -values are from one-sided Wilcoxon signed-rank tests for positive concordance (*z >* 0). Paired *P* -values test whether OncoSpot achieves higher *z*-scores than scCube. Only realizations with defined scores for both simulators were included in the paired analysis. IQR denotes interquartile range; dashes indicate not applicable.

One-sided Wilcoxon signed-rank tests demonstrated significant positive phylogeny–space concordance in HGSOC (*P* = 4.89×10^*−*8^), CRC (*P* = 7.92×10^*−*8^), and HCC (*P* = 2.59×10^*−*9^). These results indicate that malignant subclones occupying neighboring spatial spots tend to be more closely related on the simulated evolutionary tree than expected under randomized clone– tree correspondence. The consistent positive concordance across independent realizations demonstrates that the phylogenetic organization of malignant populations is reflected in their spatial distributions rather than being restricted to individual example simulations.

To assess whether OncoSpot more strongly preserves evolutionary relationships in spatial organization, we additionally compared it with scCube across matched malignant subclone populations and phylogenies. Under an identical malignant-cell-derived pseudo-spot evaluation protocol, OncoSpot achieved higher median concordance *z*-scores than scCube in HGSOC (1.250 versus 0.707), CRC (1.335 versus 0.998), and HCC (1.697 versus 0.962; **Table 2**). One-sided paired Wilcoxon signed-rank tests supported greater concordance for OncoSpot in all three cancers (*P* = 3.00 × 10^*−*4^, 8.55 × 10^*−*5^, and 1.63 × 10^*−*7^, respectively). Median paired differences remained positive across all five tested pseudo-spot grid offsets. Both simulators exhibited significant positive phylogeny–space concordance in the matched analysis, although OncoSpot consistently achieved higher median *z*-scores and stronger paired concordance across all three cancer types. This is expected, because OncoSpot explicitly incorporates the clonal phylogeny into the spatial arrangement of subclones, whereas scCube has no notion of phylogenetic relationships.

The representative HCC example in **Fig. 3** illustrates this relationship visually. The retained phylogeny contains five observed malignant subclones. Clone 0 (blue) is an ancestral population from which clone 7 (green), clone 2 (orange), clone 10 (red) descended from. Clone 21 (purple) then further descended from clone 2. In the corresponding spatial map, the three child clones (clone 7, clone 2, clone 10) are adjacent to their parent clone (clone 0). Further, clone 21 is adjacent to its parent clone clone 2. This example visually showed OncoSpot’s consistency between the phylogenetic tree and the spatial organization of malignant subclones.

**Figure 3:**
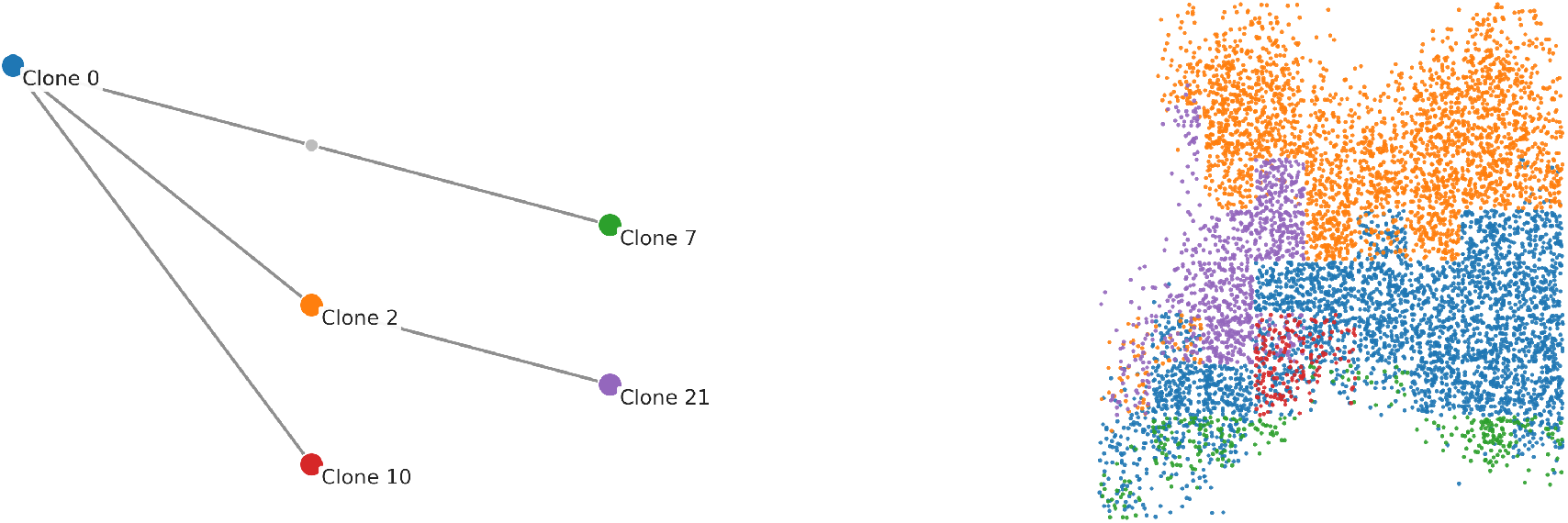
Representative HCC example illustrating phylogeny–space concordance. The left panel shows the retained malignant subclone phylogeny, with colored nodes representing observed malignant subclones and the gray node representing an ancestral-only lineage. The right panel shows the corresponding spatial distribution of individual malignant cells, colored by subclone identity. Colors are matched between the phylogeny and the spatial map. This example provides a qualitative illustration of the relationship between evolutionary ancestry and spatial organization, whereas the concordance statistics reported below were evaluated separately at the Visium spot level across independent simulations.

### Malignant spot identification across cancer types

We evaluated malignant spot identification using CopyKAT [11], CancerFinder [12], and Clonalscope [10] on the three simulated cancer datasets. Predicted malignant status was compared with the ground-truth classification derived from the cellular composition of each spot, as described in Methods. Of these three methods, CopyKAT and CancerFinder rely on transcriptomic information alone, whereas Clonalscope integrates transcriptomic and spatial information. We also considered including other state-of-the-art methods that use spatial information, such as CalicoST. However, CalicoST requires allele-level input from phased single nucleotide polymorphisms (SNPs), which the current version of OncoSpot does not simulate. Adding allele-specific copy number alterations and the corresponding allele counts is a planned extension of OncoSpot.

Classification performance varied across cancer types and computational methods (**Table 3**). CopyKAT achieved F1 scores of 0.594, 0.938, and 0.000 for HGSOC, CRC, and HCC, respectively. CancerFinder achieved corresponding F1 scores of 0.673, 0.850, and 0.203, while Clonalscope achieved 0.058, 0.913, and 0.044. CRC received the highest malignant-spot identification performance from CopyKAT and Clonalscope. In contrast, malignant spot detection in HCC was substantially less accurate across all three methods. The higher accuracy in CRC than in HCC is consistent with known differences between the two cancers. Most CRCs are chromosomally unstable, with recurrent broad arm-level copy number alterations that produce strong, genome-wide expression shifts, the signal on which CNV-based methods such as CopyKAT and Clonalscope rely [26]. In HCC, malignant hepatocytes remain transcriptionally similar to their normal counterparts [27], and the hepatocyte transcriptome is dominated by a small set of highly expressed genes, which together reduce the contrast between malignant and normal spots. That our simulated data reproduce this cancer-type-specific difficulty supports their realism.

**Table 3:** Malignant spot identification across OncoSpot-generated cancer datasets. Spots containing at least 50% malignant cells according to OncoSpot ground truth were designated tumor spots. F1 scores were calculated only for spots receiving a defined binary tumor/normal assignment from the corresponding method. Classified spots (%) is the percentage of all simulated spots assigned such a label; undefined assignments were excluded from F1 calculations.

| Cancer | Method | F1 score | Classified spots (%) |
|---|---|---|---|
| HGSOC | CopyKAT | 0.594 | 70.6 |
|  | CancerFinder | 0.673 | 100.0 |
|  | Clonalscope | 0.058 | 99.0 |
| CRC | CopyKAT | 0.938 | 94.9 |
|  | CancerFinder | 0.850 | 100.0 |
|  | Clonalscope | 0.913 | 98.3 |
| HCC | CopyKAT | 0.000 | 96.1 |
|  | CancerFinder | 0.203 | 100.0 |
|  | Clonalscope | 0.044 | 98.3 |

Unlike CancerFinder, CopyKAT and Clonalscope did not assign malignant or normal label to every spot. CopyKAT returned a defined classification for 70.6%, 94.9%, and 96.1% of spots in HGSOC, CRC, and HCC, respectively, while Clonalscope classified 99.0%, 98.3%, and 98.3% of spots in these three cancer types, as shown in **Table 3**. Spots with missing or undefined classifications (including CopyKAT’s not.defined category) were excluded from the F1 calculations. The reported F1 scores therefore describe performance among spots for which each method returned a defined binary label.

CopyKAT’s zero F1 score in HCC is consistent with our previous observation in pancreatic ductal adenocarcinoma (PDAC), where CopyKAT inverted the tumor and normal labels [28]. To investigate the differences in Clonalscope’s malignant classification performance, we examined the cellular composition of its inferred clusters. In HGSOC, Clonalscope inferred two clusters containing 213 and 201 spots. Their mean ground-truth malignant fractions were 0.852 and 0.130, respectively. Despite this separation, the malignant-enriched cluster was labeled “Normal”, while the normal-enriched cluster was labeled “Tumor”. A similar pattern occurred in HCC. The inferred clusters contained 259 and 148 spots, with mean malignant fractions of 0.729 and 0.087, respectively. These populations also received reversed malignant assignments. In contrast, Clonalscope correctly identified the predominant malignant-enriched and normal-enriched populations in CRC, consistent with its substantially higher classification F1 score. The inferred CNA burden was higher in malignant-enriched than in normal-enriched clusters across all three cancers. The respective values were 0.02088 and 0.00160 for HGSOC, 0.02039 and 0.00319 for CRC, and 0.00892 and 0.00276 for HCC.

Inspection of Clonalscope’s malignant assignment procedure revealed that it selected the two clusters with the lowest inferred CNA burden to construct a normal-reference profile. The method subsequently assigned malignant status using cosine similarity relative to that profile. For HGSOC and HCC, only two clusters were inferred, resulting in both populations being included in the reference pool. The resulting reference profiles were more closely aligned with the malignant-enriched populations, leading to reversed tumor–normal assignments.

These findings show the advantage of OncoSpot, a cancer-aware ST simulator. With known malignant and normal labels and realistic CNV-driven expression, it can reveal failure modes in existing tools that would be hard to detect in real data, where ground truth is unavailable.

### Recovery of malignant subclone identities

We next assessed whether Clonalscope could distinguish individual malignant subclones within the simulated spatial data. The dominant ground-truth malignant subclone of each spot was compared with the inferred Clonalscope cluster assignment. As detailed in Methods, evaluation was restricted to tumor-dominated spots with a dominant malignant subclone representing at least 50% of malignant cells.

Clonalscope showed limited agreement with the ground-truth subclone identities (**Table 4**). Adjusted Rand index (ARI) values were −0.0096, −0.0073, and 0.0122 for HGSOC, CRC, and HCC, respectively. Corresponding V-measure values were 0.0077, 0.0041, and 0.0269. Homogeneity scores were similarly low, ranging from 0.0021 to 0.0155.

**Table 4:** Recovery of malignant subclone identities using Clonalscope. All evaluated spots contained at least 50% (top panel) and 80% (bottom panel) malignant cells according to ground truth, respectively. Dominance denotes the minimum fraction of malignant cells assigned to the dominant ground-truth subclone. True clones and inferred clusters are the respective numbers of distinct OncoSpot malignant subclone labels and Clonalscope cluster IDs among these identical evaluation spots. All inferred-cluster counts were calculated from the saved Clonalscope assignments restricted to the corresponding evaluated spots.

| Cancer | Dominance | Spots | True clones | Inferred clusters | ARI | V-measure | Homogeneity | Completeness |
|---|---|---|---|---|---|---|---|---|
| HGSOC | ≥ 50% | 207 | 4 | 2 | −0.0096 | 0.0077 | 0.0049 | 0.0176 |
| CRC | ≥ 50% | 191 | 5 | 2 | −0.0073 | 0.0041 | 0.0021 | 0.0720 |
| HCC | ≥ 50% | 215 | 5 | 2 | 0.0122 | 0.0269 | 0.0155 | 0.1018 |
| HGSOC | ≥ 80% | 167 | 2 | 2 | −0.0081 | 0.0051 | 0.0034 | 0.0102 |
| CRC | ≥ 80% | 107 | 5 | 2 | −0.0145 | 0.0062 | 0.0033 | 0.0584 |
| HCC | ≥ 80% | 178 | 4 | 2 | 0.0043 | 0.0071 | 0.0040 | 0.0317 |

Examination of clone-by-cluster contingency tables showed extensive merging of distinct malignant subclones. In HGSOC, 196 of 207 evaluated spots (94.7%) were assigned to a single inferred cluster, although four ground-truth malignant subclones were represented. Similarly, 190 of 191 CRC spots (99.5%) and 207 of 215 HCC spots (96.3%) were assigned to their respective predominant inferred clusters, despite the presence of five ground-truth subclones in each evaluation subset.

Restricting the analysis to spots whose dominant subclone accounted for at least 80% of malignant cells did not substantially improve agreement. ARI values remained close to zero for HGSOC, CRC, and HCC. Thus, Clonalscope primarily captured broad population-level structure rather than the individual malignant subclone identities represented in these datasets. **Table 4** additionally contrasts the number of distinct true malignant subclone labels with the number of distinct Clonalscope cluster labels observed among the same evaluated tumor-dominated spots. These inferred cluster counts refer to unsupervised clone-group assignments, not Clonalscope’s subsequent binary tumor/normal labels.

### Copy-number inference using Clonalscope and CopyKAT

We evaluated copy-number inference from the simulated spatial transcriptomics data using Clonalscope [10] and CopyKAT [11]. Both methods were compared with OncoSpot’s known spot-level absolute total copy numbers (Eq. (17)). Because Clonalscope provides relative genomic-region coverage and CopyKAT provides normalized continuous CNA-associated signals, the common signal-based evaluation uses deviations from the diploid reference In more detail, Clonalscope’s diploid-calibrated estimate 2*R*_*v,r*(*g*)_ (Eq. (18)) is centered at two, whereas CopyKAT is assessed using its direct output. The matched evaluation metrics are Pearson correlation, Spearman correlation, gain-versus-loss AUROC, and signal-direction agreement, as defined in Methods.

As shown in **Table 5**, both CopyKAT and Clonalscope’s agreement with the ground truth declined from HGSOC to CRC to HCC, consistent with the decreasing copy-number burden across the three simulated cancers. Clonalscope outperformed CopyKAT in HGSOC on all four metrics (Pearson correlation, 0.505 vs. 0.422; gain-versus-loss AUROC, 0.890 vs. 0.838; directional agreement, 81.2% vs. 76.3%) and in CRC on three of four (Spearman correlation, 0.341 vs. 0.299; AUROC, 0.805 vs. 0.781; directional agreement, 74.1% vs. 68.2%), with a comparable Pearson correlation (0.261 vs. 0.268). In HCC, by contrast, CopyKAT outperformed Clonalscope on all four metrics (e.g., Pearson correlation, 0.163 vs. 0.108; AUROC, 0.714 vs. 0.692). Because the two methods report CNA signals for different sets of spots and genes, the numbers of evaluated spots and genes are given in **Table 5**.

**Table 5:** Copy-number signal concordance and profile availability for Clonalscope and Copy-KAT in tumor-dominated spatial spots. The number of spots and genes used for the reported signal metrics reflects the available observations of each method; tumor-dominated status is defined by OncoSpot ground truth (malignant-cell fraction ≥ 50%). Profile availability is the percentage of *all* simulated spatial spots for which a method returned a copy-number profile and is distinct from malignant/normal classification availability. Pearson and Spearman correlations use method-specific CNA signals relative to ground-truth *CN* − 2. AUROC distinguishes ground-truth gains (*CN* − 2 ≥ 0.5) from losses (*CN* −2 ≤ 0.5). Direction agreement compares the sign of the predicted CNA signal with the sign of the ground-truth deviation among altered gene–spot observations. Clonalscope signal is recentered by aligning the diploid-calibrated estimate at 2, whereas CopyKAT is evaluated using its native continuous signal. Clonalscope signal metrics use the full saved gene-to-region mapping, distinct from the restricted mapped genes used for its root mean squared error (RMSE) analysis.

| Cancer | Method | Spots | Genes | Profile availability (%) | Pearson | Spearman | Gain/loss AUROC | Direction (%) |
|---|---|---|---|---|---|---|---|---|
| HGSOC | Clonalscope | 216 | 71,269 | 99.0 | 0.5052 | 0.4885 | 0.8896 | 81.2 |
|  | CopyKAT | 100 | 22,761 | 70.6 | 0.4223 | 0.4039 | 0.8383 | 76.3 |
| CRC | Clonalscope | 206 | 68,190 | 98.3 | 0.2605 | 0.3413 | 0.8048 | 74.1 |
|  | CopyKAT | 200 | 19,231 | 94.9 | 0.2678 | 0.2991 | 0.7814 | 68.2 |
| HCC | Clonalscope | 218 | 68,190 | 98.3 | 0.1076 | 0.1326 | 0.6922 | 62.0 |
|  | CopyKAT | 213 | 17,099 | 96.1 | 0.1625 | 0.1668 | 0.7144 | 62.9 |

Because the method-specific gene and spot sets differ, we also compared both approaches on exactly matched gene–spot observations. The shared tumor-dominated subsets comprised 99 spots and 22,012 genes for HGSOC, 200 spots and 17,947 genes for CRC, and 211 spots and 15,992 genes for HCC. On these identical observations, Clonalscope and CopyKAT achieved Pearson correlations of 0.4789 versus 0.4300 in HGSOC, 0.2654 versus 0.2500 in CRC, and 0.0980 versus 0.1609 in HCC. Corresponding gain-versus-loss AUROC values were 0.8673 versus 0.8443, 0.8030 versus 0.7662, and 0.6946 versus 0.7159. Clonalscope performed better on all four signal-based metrics in HGSOC and CRC in this matched-observation analysis, whereas CopyKAT performed better on all four in HCC.

This cancer-type dependence is consistent with differences in the copy number landscapes of the three cancers and in the modeling assumptions of the two methods. HGSOC and CRC are characterized by extensive chromosomal instability [29, 26]. Under these conditions, Clonalscope’s segment-level modeling and use of spatial information can pool signal across large, spatially coherent genomic and tissue regions, improving CNV signal recovery. In HCC, where the simulated phylogenies have shorter edges and therefore fewer alterations, the copy number signal is weaker and less extensive [27]. Strong segment-level and spatial smoothing may then merge or miss the fewer events, whereas CopyKAT’s gene-level smoothing may retain these weaker shifts more effectively. Because such method-by-cancer interactions and comparisons cannot be effectively quantified in real tumors as they often lack ground truth, OncoSpot being a cancer-aware simulator provides realistic, cancer-type-specific data with known copy number profiles, and has the potential to become a principled basis for selecting the most suitable method for a given cancer type.

Since Clonalscope provides the absolute copy number for each gene at each spot, we further assessed the quantitative error of its diploid-calibrated total-copy-number estimates (Eq. (18)) against OncoSpot’s cell-averaged absolute total-copy-number ground truth (Eq. (17)). A constant-diploid estimate was evaluated over the same matched gene–spot observations as a baseline. In tumor-dominated spots, Clonalscope achieved copy-number root mean squared error (RMSE) values of 0.6772, 0.8701, and 0.6169 for HGSOC, CRC, and HCC, respectively (**Table 6**). The corresponding constant-diploid baseline RMSE values were 0.8029, 0.8964, and 0.5399. Relative to the diploid baseline, Clonalscope reduced RMSE by 15.7% in HGSOC and 2.9% in CRC. In contrast, its RMSE was 14.3% higher than the baseline in HCC. That Clonalscope’s accuracy decreases in HCC is consistent with the observation in **Table 5**. We further examined gene–spot pairs whose true total copy number differed from diploid by at least 0.5. Within this subset, Clonalscope achieved RMSE values of 0.9816, 1.3705, and 1.0988 for HGSOC, CRC, and HCC, respectively. The corresponding diploid-baseline errors were 1.2611, 1.4929, and 1.1192, indicating that Clonalscope estimates absolute copy number more accurately in altered regions than in copy-neutral regions. Such region-specific assessment is difficult with real data, where ground-truth copy number is rarely available at spot resolution, but is straightforward with OncoSpot, which explicitly models cancer CNAs and clonal structure.

**Table 6:**
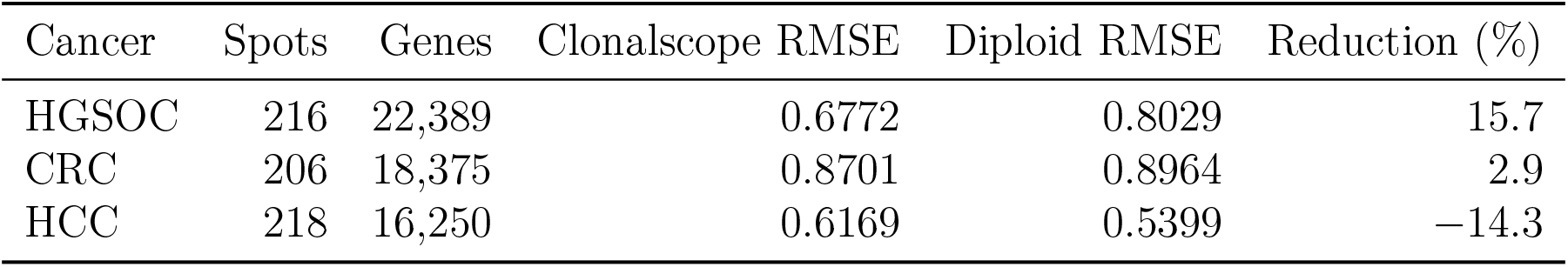
Clonalscope diploid-calibrated total copy-number reconstruction in tumor-dominated spots. RMSE was calculated against OncoSpot’s cell-averaged absolute total-copy-number ground truth using Eqs. (20) and (21). The same matched genes and spots were used for the Clonalscope and diploid baseline calculations. Positive RMSE reduction indicates improvement over the baseline.

### Cell-type deconvolution using RCTD

We further evaluated whether OncoSpot-generated spatial transcriptomics data could support benchmarking reference-based cell-type deconvolution. We applied RCTD (spacexr v2.2.1) in full mode to the simulated HGSOC, CRC, and HCC datasets, using empirical single-cell RNA-sequencing references previously employed for cancer-specific OncoSpot calibration.

The reference cells were grouped into seven broad categories: malignant, T cells, B/plasma cells, myeloid/macrophage cells, fibroblasts, endothelial cells, and normal epithelial cells. All simulated spatial spots were provided to RCTD, but quantitative evaluation was restricted to purely nonmalignant spots that contain no malignant cells according to the OncoSpot ground truth. This is because RCTD is not designed to deconvolve spots containing malignant cells.Predicted compositions of each of the six nonmalignant cell types were compared with the corresponding ground-truth cell fractions using RMSE across spots and cell types and mean Jensen–Shannon divergence (JSD; logarithm base 2) across spots. RCTD produced valid predictions for all 329 purely nonmalignant spots across the three cancer types (104 in HGSOC, 125 in CRC and 100 in HCC; **Table 7**). Deconvolution accuracy was broadly comparable across cancer types, with RMSEs of 0.131, 0.134 and 0.143 and mean Jensen–Shannon divergences (JSDs) of 0.088, 0.083 and 0.101 in HGSOC, CRC and HCC, respectively. Errors were lowest in HGSOC and CRC, and slightly higher in HCC on both metrics.

**Table 7:** RCTD cell-type deconvolution performance on pure-normal spatial spots. RCTD was applied to all simulated spots, but quantitative evaluation was restricted to spots containing no malignant cells according to OncoSpot ground truth. Predicted weights for the six nonmalignant cell types were normalized to sum to one before comparison with ground-truth cell fractions. RMSE was calculated across all evaluated spot–cell-type combinations, and mean JSD was calculated across spots using logarithm base 2. All pure-normal spots received valid RCTD predictions. Lower RMSE and JSD indicate better agreement with the simulated cell-type compositions.

| Cancer | Pure-normal spots | Tumor spots excluded | RMSE | Mean JSD |
|---|---|---|---|---|
| HGSOC | 104 | 314 | 0.1308 | 0.0879 |
| CRC | 125 | 284 | 0.1336 | 0.0828 |
| HCC | 100 | 314 | 0.1434 | 0.1014 |

**Fig. 4** compares the true and RCTD-inferred spatial distributions of nonmalignant cell-type compositions. The predicted compositions broadly reproduce the spatial variation in cell-type mixtures represented in the ground truth, although differences in individual cell-type proportions remain visible. Tumor-containing spots are displayed for spatial context but excluded from quantitative evaluation. Together, these results demonstrate the feasibility of benchmarking reference-based cell-type deconvolution to OncoSpot-generated data across distinct cancer-specific simulation configurations.

**Figure 4:**
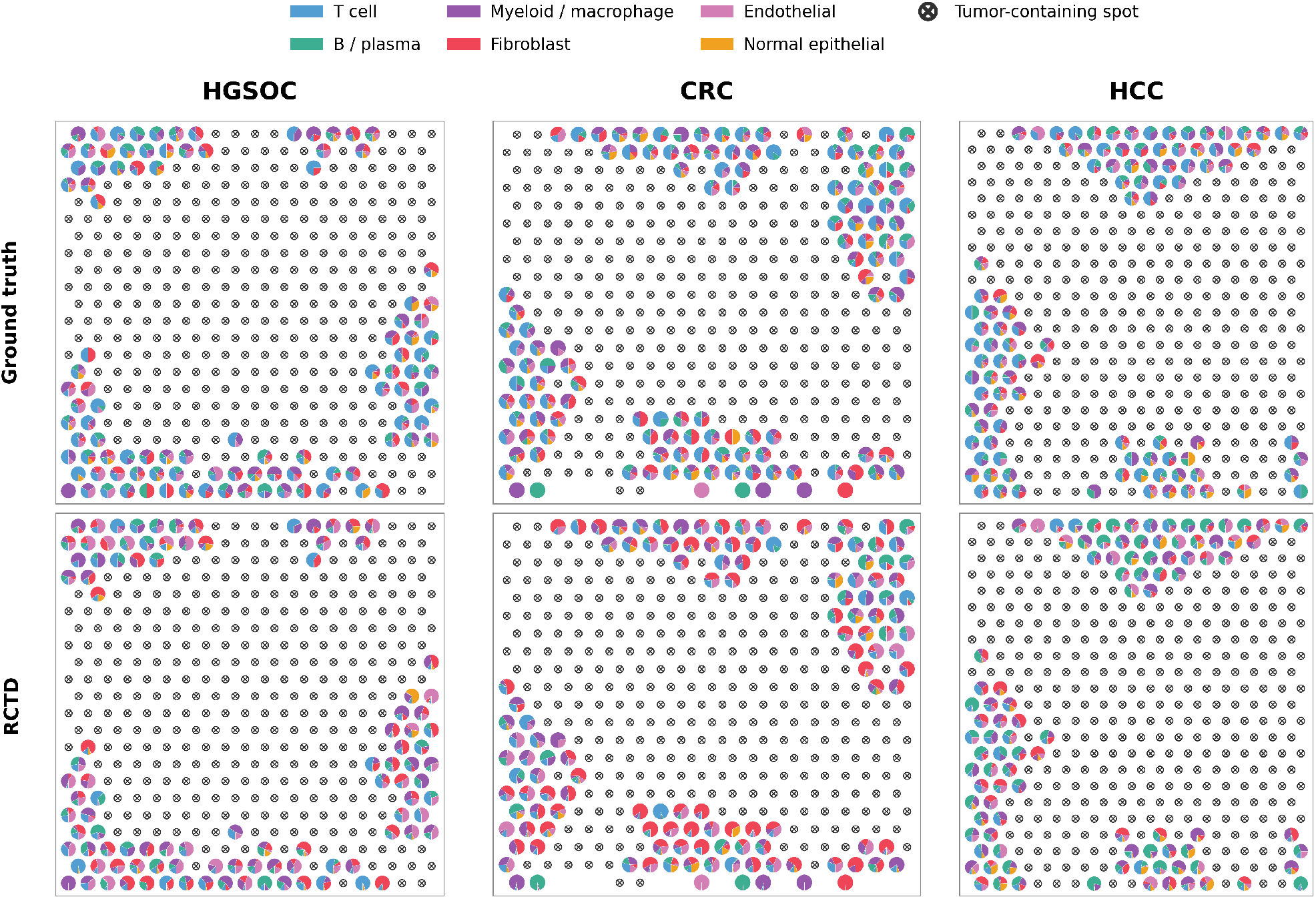
Comparison of ground-truth and RCTD-inferred nonmalignant cell-type compositions in OncoSpot-simulated spatial transcriptomics data. The columns represent high-grade serous ovarian carcinoma (HGSOC), colorectal cancer (CRC), and hepatocellular carcinoma (HCC). The top row shows ground-truth cell-type compositions, while the bottom row shows RCTD-inferred compositions at the corresponding spatial locations. Each pie chart represents the relative proportions of six nonmalignant cell types within a pure-normal spot. Spots containing any malignant cells are indicated by crossed-circle markers in both rows and were excluded from quantitative evaluation.

## Discussion

We used HGSOC, CRC and HCC to represent three of the four evolutionary modes described by Noble et al. [23]: branching evolution (HGSOC; invasive glandular growth), progressive diversification (CRC; gland fission), and effectively almost neutral evolution (HCC; boundary growth). The fourth, non-spatial mode (selective sweeps), represents hematologic malignancies such as leukemia, which fall outside the current scope of OncoSpot, as discussed below.

OncoSpot is nevertheless not restricted to the three cancer types analyzed in this study. Its modular design separates the evolutionary scaffold from the cancer-specific components, so users can match the spatial growth architecture to the cancer being simulated by selecting among the growth mechanisms that OncoSpot exposes through its DEMON interface. This choice also shapes the copy number landscape: OncoSpot’s CNA generation procedure is reused across cancer types, but the number of alterations along each lineage is derived from the edge lengths of the selected scaffold’s phylogeny, so the CNA burden automatically reflects the evolutionary mode of the chosen cancer type. Cancer-specific transcriptomic reference data can then be supplied, while OncoSpot’s downstream framework for cell-level expression, spatial compartments and spot-level observation is reused unchanged. For example, breast cancer can be simulated by selecting the invasive glandular growth mode, which Noble et al. associate with breast cancer [23], together with breast-specific expression references.

To help users apply this strategy, we reviewed evidence on spatial growth and clonal evolution across additional malignancies and identified the closest DEMON growth mode for several representative cancer contexts (**Table 8**). The mapping is based primarily on tumor growth architecture rather than on cancer name alone. Infiltrative solid tumors are most naturally represented by invasive glandular growth, whereas tumors with predominantly pushing or expansile borders may be better represented by boundary growth. Gland fission is most directly supported for early colorectal glandular lesions [23]. Some malignancies, including sarcomas and neuroendocrine tumors, show heterogeneous growth architectures and therefore do not support a single cancer-wide assignment. Because OncoSpot is implemented as an end-to-end pipeline, these choices require no direct interaction with DEMON, which comes with a wrapper interface that lets users specify the cancer type or growth mode.

**Table 8:**
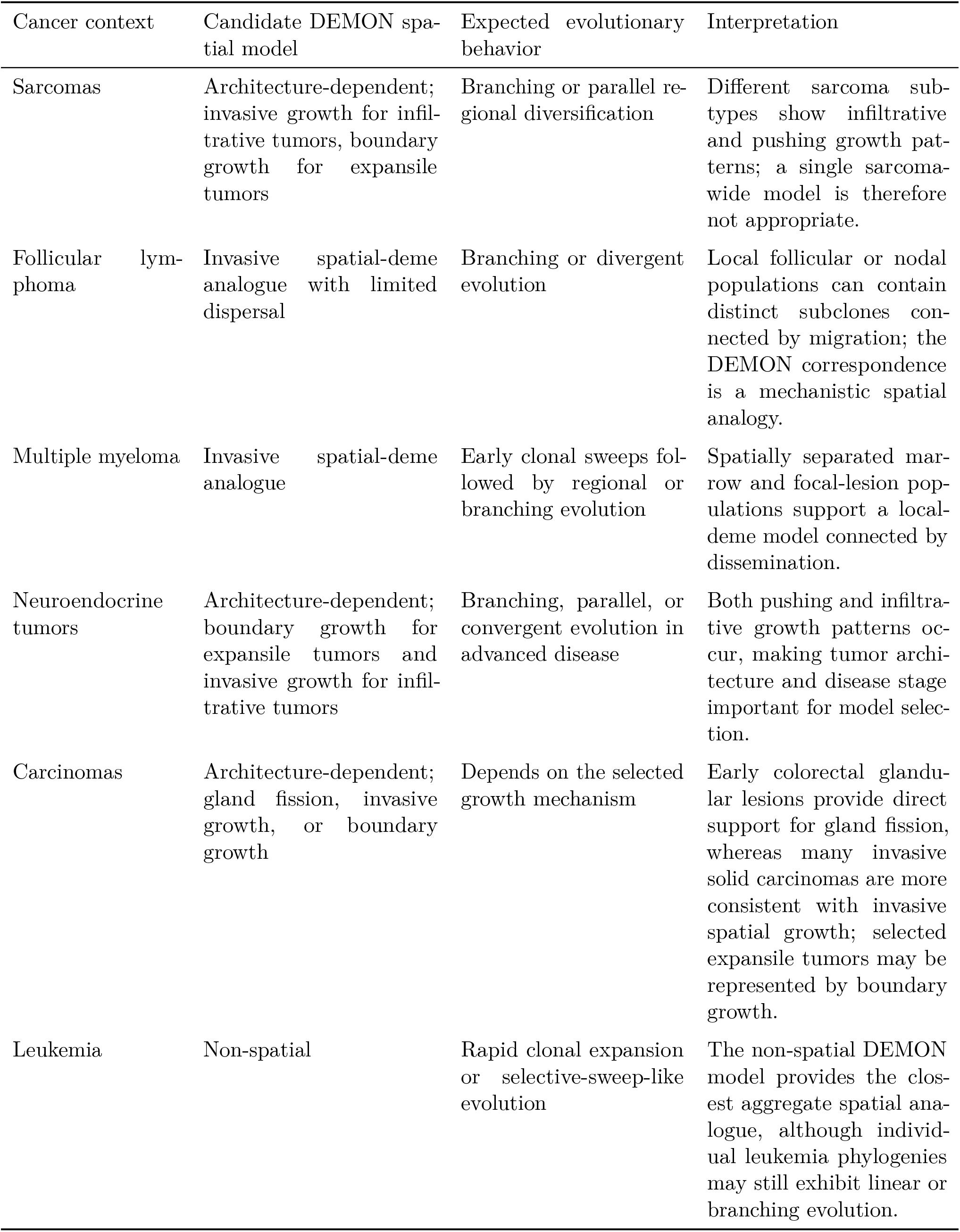
Potential extension of OncoSpot to additional cancer contexts using alternative DEMON spatial growth models. Assignments describe the closest mechanistic analogue supported by tumor architecture and evolutionary evidence and should not be interpreted as universal models for all tumors within a cancer category.

| Cancer context | Candidate DEMON spa-tial model | Expected evolutionary behavior | Interpretation |
|---|---|---|---|
| Sarcomas | Architecture-dependent; invasive growth for infil-trative tumors, boundary growth for expansile tumors | Branching or parallel re-gional diversification | Different sarcoma sub-types show infiltrative and pushing growth pat-terns; a single sarcoma-wide model is therefore not appropriate. |
| Follicular lym-phoma | Invasive spatial-deme analogue with limited dispersal | Branching or divergent evolution | Local follicular or nodal populations can contain distinct subclones con-nected by migration; the DEMON correspondence is a mechanistic spatial analogy. |
| Multiple myeloma | Invasive spatial-deme analogue | Early clonal sweeps fol-lowed by regional or branching evolution | Spatially separated mar-row and focal-lesion pop-ulations support a local-deme model connected by dissemination. |
| Neuroendocrine tumors | Architecture-dependent; boundary growth for expansile tumors and invasive growth for infil-trative tumors | Branching, parallel, or convergent evolution in advanced disease | Both pushing and infiltra-tive growth patterns oc-cur, making tumor archi-tecture and disease stage important for model selec-tion. |
| Carcinomas | Architecture-dependent; gland fission, invasive growth, or boundary growth | Depends on the selected growth mechanism | Early colorectal glandu-lar lesions provide direct support for gland fission, whereas many invasive solid carcinomas are more consistent with invasive spatial growth; selected expansile tumors may be represented by boundary growth. |
| Leukemia | Non-spatial | Rapid clonal expansion or selective-sweep-like evolution | The non-spatial DEMON model provides the clos-est aggregate spatial ana-logue, although individ-ual leukemia phylogenies may still exhibit linear or branching evolution. |

OncoSpot is designed to simulate spot-based Visium measurements of sectioned solid tumors, and hematologic malignancies, including circulating blood cancers and bone-marrow-resident disease, are therefore not included. Circulating malignant cells do not form the persistent, sectioned tissue geometry assumed by the deme scaffold and the spot-capture model. Bone marrow, in contrast, is a spatially organized tissue that can be assayed by spatial transcriptomics. Its exclusion reflects differences in tissue organization and specimen processing rather than an absence of spatial structure. Decalcification of bone-containing biopsy material can compromise RNA and morphological preservation, while adipose-rich marrow regions, variable cellular density, and autofluorescence in imaging workflows introduce additional tissue-dependent sampling and quality-control considerations [30, 31]. Successful spatial profiling of marrow has nevertheless been reported, including with Visium, so these obstacles are not absolute platform restrictions [30]. Modeling marrow would, however, require a scaffold and observation model tailored to this tissue rather than the solid-tumor deme geometry used here.

We focused this first version of OncoSpot on Visium because it is among the most widely used platforms for profiling tumor sections and because its transcriptome-wide capture provides the genome-wide coverage on which expression-based CNA inference depends, which is central to the benchmarks presented here. Imaging-based platforms such as Xenium differ from Visium in two respects that prevent a direct extension. First, they measure targeted gene panels whose genomic coverage is far sparser than transcriptome-wide capture, which limits the applicability of current CNA inference methods. Second, they require a fundamentally different observation model: rather than aggregating transcripts within capture spots, they resolve individual RNA molecules, so simulating them requires modeling cell and nuclear boundaries, three-dimensional section geometry or optical sampling, transcript detection and decoding, and cell segmentation [32]. Supporting these modalities therefore goes beyond reassigning Visium spot coordinates, and we leave it to future work.

## Methods

### Computational framework and mathematical model

#### Overview and notation

OncoSpot generates a spatially organized population of malignant and nonmalignant cells and a Visium-like, spot-resolution transcriptomic observation from a simulated tissue section. The observation model is restricted to spot-based spatial transcriptomics rather than imaging-based, single-molecule assays. Its components are linked in the following order: deme-based tumor evolution with DEMON [23], construction of a spatial cell scaffold and tissue compartments, copy-number evolution on the malignant subclone phylogeny with CNAsim [24], cell-level expression sampling, and aggregation into Visium-like spots. This division separates the scale at which evolution is simulated (demes and genetic lineages), the scale at which RNA counts are simulated (individual cells), and the scale at which spatial RNA counts are observed (i.e., spots).

Let *i* = 1, …, *N* index simulated cells, *g* ∈ G index genes, *c*(*i*) denote the malignant subclone of malignant cell *i*, and *t*(*i*) denote the broad cell type of nonmalignant cell *i*. The spatial coordinate of a cell is **r**_*i*_ = (*x*_*i*_, *y*_*i*_). Each malignant cell has a state-activity vector **z**_*i*_ = (*z*_*i*1_, …, *z*_*iK*_)^*⊤*^, where *K* is the number of cancer-associated transcriptional states modeled (e.g., proliferation, hypoxia, epithelial–mesenchymal transition, and stress response), and *z*_*ik*_ denotes the activity of state *k* in cell *i*. We denote by *CN*_*gc*_ the total copy number of gene *g* inherited by malignant subclone *c*, and by *L*_*i*_ the positive library-size scale sampled for cell *i*. In the following, we describe in detail each component of the model. **Supplemental Section S2** lists the model and simulation parameters.

#### Deme-based malignant evolution and spatial scaffold

Malignant evolutionary histories are generated with DEMON, a deme-based population-genetic simulator that models birth, death, driver and passenger mutation, selection, and cell dispersal under alternative tissue architectures [23]. We use cancer-specific configurations to represent distinct mechanistic growth scenarios rather than imposing a fixed number of subclones. The mapping of a cancer type to a DEMON architecture is supported by tissue morphology and evolutionary observations to differing degrees, and is not assumed to describe every tumor of that type.

For high-grade serous ovarian carcinoma (HGSOC), we select the *invasive-glandular* DEMON configuration as an analogue of invasion by spatially localized malignant populations. Histopathological descriptions include glandular or papillary components, destructive stromal invasion, and marked variation in tissue architecture [33, 34]. Multiregion phylogenetic studies further show spatially heterogeneous subclones and divergent intraperitoneal dissemination [35]. These observations motivate an invasive, spatially structured scenario.

For colorectal cancer (CRC), we adopt *gland fission* to model an early glandular growth scenario. Human colorectal adenomas have direct histological evidence of crypt expansion through fission [36], and DEMON explicitly implements this architecture as a model of colorectal adenoma growth [23]. This is a model of one CRC-related growth mechanism, not an assertion that advanced invasive colorectal carcinomas continue to evolve exclusively through gland fission; a transition to invasive growth is biologically plausible [23].

For hepatocellular carcinoma (HCC), we adopt *boundary-restricted growth* as an analogue of expansile tumors whose proliferation and outward expansion are concentrated near the tumor boundary. Histopathological work distinguishes expansile or encapsulated HCC from replacing and sinusoidal/infiltrative patterns [37, 38]. Noble et al. specifically discuss boundary-growth models for selected HCC contexts while emphasizing heterogeneous HCC architectures [23]. The literature basis and limits of these correspondences are summarized alongside the scenario parameters in **Supplemental Section S1**.

Each DEMON realization provides deme locations, deme-specific cell abundances, driver-genotype assignments, and a driver phylogeny; thus, the number of cells associated with each DEMON deme is determined by the DEMON simulation rather than fixed a priori. To obtain cell-level spatial coordinates, the resulting cell records are redistributed over the filled spatial scaffold while preserving their DEMON lineage composition, and each cell is assigned a continuous position by random jitter within its corresponding spatial tile.

#### Malignant compartment and spatial microenvironment

The DEMON-generated cell population provides the spatial scaffold for tissue construction, but its internal normal-cell annotations are not used as the final tumor-microenvironment assignments. This is because DEMON labels the entire tumor mass as malignant and confines normal cells to the outer margin, so it cannot represent infiltrating normal cells. To address this issue, starting from an unlabeled cell population, we assign each cell a malignant or nonmalignant identity in successive steps, aiming to produce a malignant-enriched tissue interior and a nonmalignant-enriched periphery while allowing irregular boundaries and spatial mixing. We then assigned the cell type to nonmalignant cells afterwards. Following paragraphs give more details on this whole process.

For every position (*u, v*) of a cell *i*, we determine whether it will be a nonmalignant score based on its nonmalignant score *Q*_*i*_. *Q*_*i*_ will be a combination among a boundary field *B*(*u, v*) which is higher when (*u, v*) is closer to unlabeled cells, a smoothed normalized random value drawn from a Gaussian distribution *R*(*u, v*), and a random value to introduce cell-level randomness *ϵ*_*i*_. Here we describe these three terms, respectively.

Starting from (*u, v*), a position of a cell on the two-dimensional space, let *D*(*u, v*) denote its Euclidean distance to the nearest unlabeled location, and *D*_max_ be the largest such distance across the labeled cells.

We define the boundary field, *B*(*u, v*), as

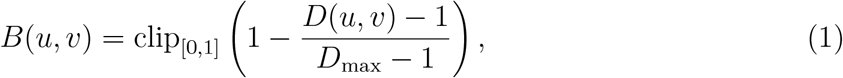

for *D*_max_ *>* 1. The function clip_[0,1]_ restricts values to the interval [0, 1]. The closer (*u, v*) is to the boundary of the assigned cells, the higher *B*(*u, v*) is.

Because real tumor–stroma interfaces are spatially irregular, boundary proximity alone is insufficient to determine compartment assignment. We therefore introduce a spatially correlated random field that permits nonmalignant regions to extend into the tissue interior. Independent standard-normal random values *ϵ*_*R*_(*u, v*) are generated on the deme grid and smoothed using a Gaussian kernel:

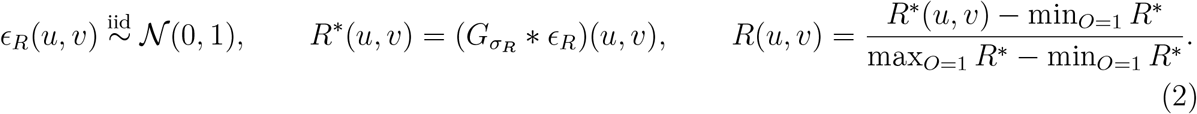

Here, 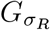 denotes a Gaussian smoothing kernel with spatial scale *σ*_*R*_, * denotes convolution, and the minimum and maximum are calculated over occupied tissue locations. The normalized field *R*(*u, v*) varies between zero and one, producing smoothly varying spatial preferences rather than independent random assignments at each location.

Finally, we combine the boundary field and the spatial random field to determine which cells belong to the nonmalignant compartment. For cell *i*, let *B*_*i*_ and *R*_*i*_ denote the corresponding field values at its spatial location. A nonmalignant score of cell *i* is calculated as

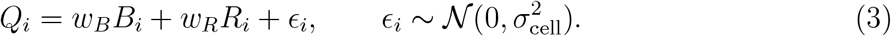

Here, *w*_*B*_ controls the preference for peripheral placement, *w*_*R*_ controls the contribution of spatially correlated variation, and *ϵ*_*i*_ introduces cell-level randomness with standard deviation *σ*_cell_.

Given a pre-determined number of nonmalignant cells, *N*_*normal*_, which can be calculated from the total number of cells and a user-given nonmalignant cell proportion, we rank the nonmalignant score of all cells, and the top *N*_*normal*_ cells with the highest nonmalignant score will be assigned as nonmalignant cells. This procedure preserves the specified malignant and nonmalignant cell counts exactly while favoring a malignant-enriched interior and a spatially heterogeneous nonmalignant periphery.

After malignant and nonmalignant cells are labeled, we proceed to define the cell type of each nonmalignant cell. Nonmalignant cells are annotated using broad T-cell, myeloid/macrophage, fibroblast, endothelial, B/plasma, and normal-epithelial categories. Because nonmalignant cells are not randomly distributed in tissue, we assign their cell types with spatial dependence, so that cells of the same type are more likely to be neighbors. For nonmalignant cell *i*, we define its distance to the nearest malignant cell, *d*_*i*_ as *d*_*i*_ = min_*j∈ℳ*_ ∥**r**_*i*_ − **r**_*j*_∥_2_, where *ℳ* denote malignant cell sets. We also calculate the median of its distance to the *k*-nearest nonmalignant cells, *σ*_*T*_. We define the tumor-interface score for cell *i, T*_*i*_, as

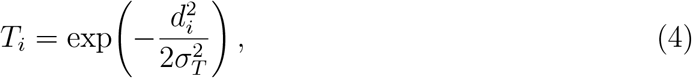

The nonmalignant cells are linked by a *k*-nearest-neighbor graph. During sequential labeling, the relative assignment weight for cell type *t* is

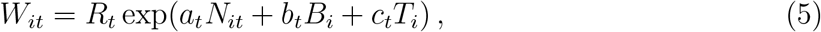

where *R*_*t*_ denotes the remaining number of cells required for type *t, N*_*it*_ is the fraction of already assigned neighbors belonging to that type, and (*a*_*t*_, *b*_*t*_, *c*_*t*_) quantify neighborhood, peripheral, and tumor-interface preferences. Label propagation starts near the malignant– nonmalignant interface and preserves the specified overall counts for each cell type. The factor *R*_*t*_ is updated after each assignment and becomes zero once the prescribed number of cells of type *t* has been reached, preventing further assignments to that type. Thus, positive *a*_*t*_ encourages spatial clustering among cells of the same type, while *R*_*t*_ ensures that the target cell-type abundances are preserved.

#### Subclone-aware CNA simulation and gene-level mapping

After tissue compartments have been assigned, malignant subclone abundances are calculated from the final malignant-cell population. To avoid treating extremely rare evolutionary lineages as separately resolvable benchmark clones, subclones representing less than 1% of malignant cells are collapsed into the nearest retained subclone on the DEMON driver phylogeny. Phylogenetic proximity is defined by the shortest unweighted path between clone nodes, with the ties broken by an ancestral clone followed by the more abundant retained clone.

The DEMON driver phylogeny is then restricted to the retained malignant subclones and any ancestral nodes required to connect them, with irrelevant branches pruned. The resulting malignant subclone tree is supplied to CNAsim [24]. CNAsim simulates copy-number alterations along the retained phylogeny. The resulting segment-level total-copy-number profiles are mapped to genomic genes, yielding *CN*_*gc*_ ∈ {0, 1, 2, …} for each retained clone *c* and covered gene *g*. Every malignant cell inherits the corresponding profile through its final subclone ID. CNAs are not resampled independently for every cell or deme. Adding cell-specific CNAs will be in the future work. Nonmalignant cells are modeled without malignant clone-specific CNA evolution and have a diploid reference copy number for expression and CNA ground-truth annotation. Original DEMON assignments, clone-collapse mappings, clone-level CNA profiles, malignant-cell-to-clone assignments, gene mappings, and the retained subclone phylogeny are preserved as ground truth and provenance.

#### Reference-based expression for normal cells

For each cancer type, a set of real datasets is used to estimate the baseline transcript composition for the normal cells. Specifically, we use early-secretory fallopian-tube epithelial cells for HGSOC, normal colonic stem-like/transit-amplifying epithelial cells for CRC, and normal-site hepatocytes for HCC’s baseline transcript. To prevent a patient contributing many reference cells from dominating the baseline, counts are first aggregated within each patient. Let 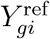 denote the observed count of gene *g* in reference cell. Let 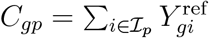 be the pseudobulk count of gene *g* in donor *p*. Define

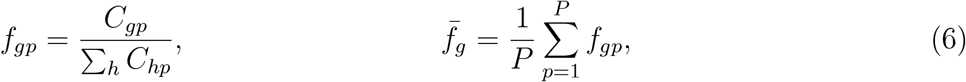

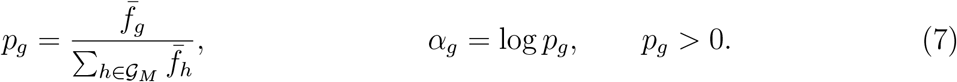

Thus *p*_*g*_ is a relative baseline fraction and *α*_*g*_ is its log value. Only positive-baseline genes are included in the malignant expression branch. Patients contribute equally to the baseline, irrespective of reference cell counts. Baselines are calculated from reference counts rather than corrected integrated expression values.

#### Empirical malignant state activities and gene-level effects

Beside the CNAs, we add malignant state activities stochastically to reflect the malignant transcriptional heterogeneity of each cancer type. Six common Hallmark-derived activity axes in HGSOC, CRC, and HCC are used: proliferation (the union of E2F targets and G2M checkpoint), WNT/*β*-catenin signaling, MYC targets, hypoxia, epithelial–mesenchymal transition (EMT), and glycolysis (denoted METABOLIC) [39]. These continuous axes are potentially overlapping transcriptional programs rather than mutually exclusive cell states. Nevertheless, together they span the major recurrent sources of malignant transcriptional heterogeneity, namely cell-cycle activity, oncogenic signaling (WNT/*β*-catenin and MYC), response to the microenvironment (hypoxia), cellular plasticity (EMT), and metabolic reprogramming (glycolysis), several of which also emerge as recurrent intratumoral programs across cancer types [8]. We calibrate their distributions and gene-level associations separately for each cancer because both the prevalence of these programs and the genes through which they are expressed differ between tumors of different tissue origins. To preserve within-cell covariation among programs, OncoSpot does not use the DEMON phylogeny for transcriptional states assignment. Instead, we resample the entire reference-cell vectors, 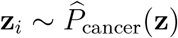.

For gene *g* at cell *i*, the log-normalized reference expression *x*_*gi*_ is 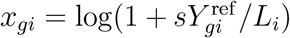, where *s* denotes the normalization scale and *L*_*i*_ is the reference-cell UMI library size. Genes with negligible across-cell variation are discarded, and retained signature genes are standardized gene-wise across reference cells. For each program, standardized values are averaged across its usable signature genes and the resulting score is centered and standardized to unit population variance. Thus, the six state coordinates are on comparable scales within each cancer, while their empirical joint distributions remain cancer-specific.

We define *γ*_*gk*_ as the gene-level response coefficient for gene *g* associated with Hallmark-derived activity axis *k*. These coefficients are estimated using ridge regression of log-normalized reference counts on the six state scores. Let *N*_ref_ denote the number of reference cells and 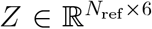 denote the reference-cell-by-state matrix, where each row corresponds to the six-dimensional state-activity vector of a reference cell. Let *X* be the patient-centered version of *Z*, obtained by subtracting the corresponding patient-specific mean from each state score. Similarly, let **y**_*g*_ denote the vector of log-normalized expression values for gene *g* across reference cells, centered within each patient. The gene-specific ridge regression coefficients are estimated as

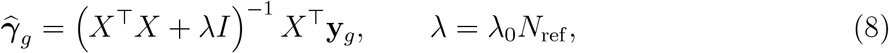

Here, 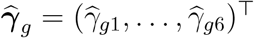 contains the estimated response coefficients for gene *g* across the six transcriptional axes, *I* is the 6 × 6 identity matrix, *λ* is the ridge regularization parameter, and *λ*_0_ is a nonnegative baseline regularization constant controlling the strength of coefficient shrinkage.

Centering within the sole HGSOC reference patient is equivalent to global centering of its selected cells. Likewise, CRC and HCC reference cells are centered within their respective patients. Coefficients are limited componentwise to [−*γ*_max_, *γ*_max_], and genes below detection or mean-expression thresholds receive zero coefficients.

The combined effect of the six transcriptional activity axes on gene *g* in malignant cell *I* is defined as

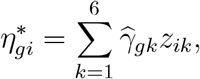

where *z*_*ik*_ is the activity of axis *k* in cell *i*. To prevent extreme transcriptional modulation, this effect is bounded using gene-specific empirical reference quantiles:

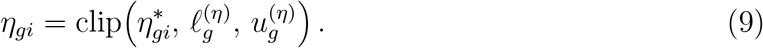

Here, 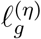 and 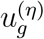 denote the gene-specific lower and upper clipping bounds, respectively, derived from the empirical reference distribution of state-induced expression effects. The clip function restricts 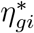 to this interval.

The estimated associations provide a regularized expression-modulation mechanism rather than a causal separation of transcriptional programs or a negative-binomial generalized linear-model fit. This is to avoid overfitting and produce robust, plausibly sized expression shifts.

#### Malignant cell-level RNA counts

Given malignant cell *i* in clone *c*(*i*) and gene *g* with *CN*_*g,c*(*i*)_ *>* 0, its conditional mean expression is

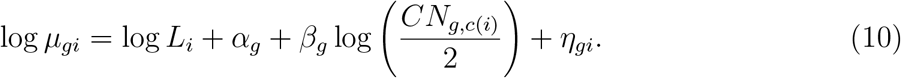

Here, *L*_*i*_ represents the cell-specific sequencing depth, *α*_*g*_ specifies the baseline log-expression fraction of gene *g, β*_*g*_ quantifies the gene-specific transcriptional response to copy-number changes, *CN*_*g,c*(*i*)_ denotes the copy number of gene *g* in the malignant clone containing cell *i*, and *η*_*gi*_ captures cell-specific transcriptional variation arising from malignant cell states and the tumor microenvironment.

Equivalently, 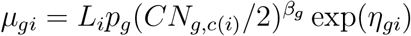. The exponent *β*_*g*_ controls the strength of the positive-copy-number dosage association, with copy number two serving as the reference. CNA-dependent transcriptional modulation is motivated by paired single-cell DNA/RNA studies [9]. For homozygous deletion (*CN*_*g,c*(*i*)_ = 0), *µ*_*gi*_ = 0 is enforced explicitly.

Counts follow the negative-binomial NB model,

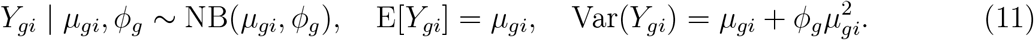

A nonnegative reference abundance is chosen so that _*i*_ *µ*_*gi*_ matches the observed tumor count total after accounting for sampled library sizes and bounded state effects, and the raw moment estimator is

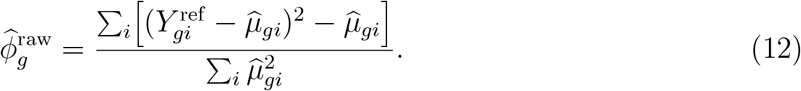

Finite estimates are clipped to [0, *ϕ*_max_]. When the moment estimate is undefined, a prespecified fallback *ϕ*_fallback_ is used.

#### Nonmalignant cell-level RNA counts

Nonmalignant expression is calibrated separately for each broad cell type. Let *p*_*gt*_ denote the type-specific baseline transcript fraction obtained from reference single-cell counts, with ∑_*g*_ *p*_*gt*_ = 1, and let *ϕ*_*t*_ denote a pooled NB dispersion for cell type *t*. The simulated expression for nonmalignant cell *i* of type *t*(*i*) is

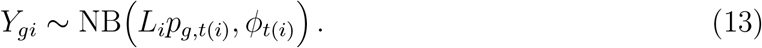

Empirical reference library-size distributions are used to sample *L*_*i*_ within the malignant and nonmalignant compartments. Nonmalignant cells do not acquire malignant clone-specific CNA dosage effects. The malignant and nonmalignant matrices are written in sparse chunks and merged over their gene-set union *G* = *G*_*M*_ ∪ *G*_*N*_, with zero counts for gene–branch combinations absent from one branch.

#### Visium-like spot observation

The latent cell population is observed with a Visium-like, sequencing-based spot-capture layer. Spot centers are placed on a regular hexagonally staggered grid. The physical spot diameter and center-to-center spacing of the reference platform are scaled to the dimensionless simulated tissue coordinates. The spatial scale is selected to achieve a configurable target number of captured cells per occupied spot. Suppose for a spot *v* whose center is **q**_*v*_, and cell *i*’s position **r**_*i*_, with effective radius *R*_*V*_, the set of captured cells is

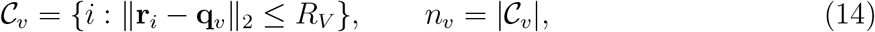

in which *n*_*v*_ is the number of cells from spot *v*. For a spot *v* that is nonempty, we calculate its latent expression for gene *g, θ*_*gv*_, as the sum across all its cells, normalized by each gene:

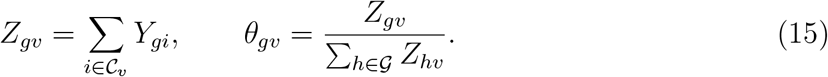

Spot depth *U*_*v*_ is sampled from an empirical Visium UMI-depth histogram derived from the GSE211956 reference, using the configured observation preset [4, 40]. The final observed spot-expression vector is

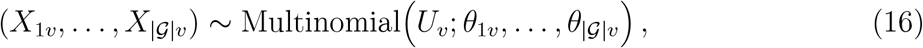

in which |G| is the total number of genes simulated. Consequently Σ_*g*_ *X*_*gv*_ = *U*_*v*_. By aggregating after cell-level RNA simulation, the observation layer represents mixtures of malignant subclones and nonmalignant cell types without assigning a single transcriptomic program to each spot. The generated spot ground truth includes captured-cell counts, malignant fractions, cell-type proportions, malignant-subclone mixture fractions, and dominant malignant subclone when present. Details of the observation model, output and a quantified configuration check are in **Supplemental Section S3**.

### Evaluation of copy-number reconstruction and malignant subclone inference

#### Spot-level absolute copy-number ground truth

OncoSpot retains the absolute total copy-number profile of each malignant subclone and the identities of all cells contributing to each simulated spatial spot. These quantities enable direct construction of the spot-level copy-number ground truth.

Let *C*_*v*_ denote the set of cells contributing to spot *v*, with *n*_*v*_ = |*C*_*v*_|. For malignant cell *i*, the copy number of gene *g* is *CN*_*g,c*(*i*)_, where *c*(*i*) denotes its malignant subclone. Nonmalignant cells are assigned the reference diploid copy number of two. We define the absolute total copy number of gene *g* in spot *v* as

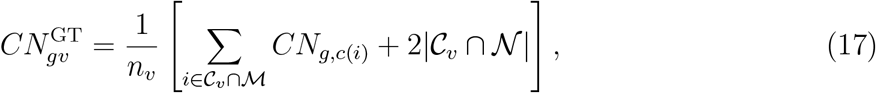

where *M* and *N* denote the malignant and nonmalignant cell sets, respectively.

This definition preserves contributions from multiple malignant subclones and nonmalignant cells within a spatial spot. It represents the cell-averaged absolute copy number rather than the copy-number profile of the dominant malignant subclone.

Ground-truth profiles were reconstructed from OncoSpot’s clone-level gene copy numbers, malignant-cell-to-clone assignments, and cell-to-spot mappings. The reconstructed cell counts and malignant-cell counts were checked against the simulation’s spot-level ground truth.

#### Copy-number reconstruction using Clonalscope

Clonalscope [10] was applied to the simulated spot-level RNA count matrices, and its inferred genomic-region coverage profiles were retained without modification.

Clonalscope reports relative coverage rather than direct absolute DNA copy-number measurements. To assess copy-number reconstruction on a diploid-referenced scale, the inferred relative coverage *R*_*vr*_ for spot *v* and genomic region *r* was converted to

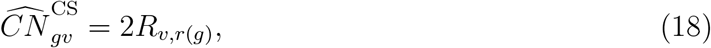

where *r*(*g*) denotes the inferred genomic interval containing the midpoint of gene *g*.

This transformation maps relative coverage of one to the diploid reference value of two. It provides a diploid-calibrated total-copy-number estimate under an assumed diploid baseline, rather than an independent estimate of absolute tumor ploidy. Genes without an assigned Clonalscope genomic interval were excluded. For signal-based comparison with CopyKAT, the diploid-calibrated Clonalscope estimates are expressed as deviations from two,

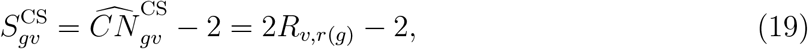

which are evaluated against the corresponding ground-truth copy-number deviations. The same Pearson correlation, Spearman correlation, gain-versus-loss AUROC, and directional-agreement definitions used for CopyKAT are applied to these centered Clonalscope signals; no refitting of the underlying Clonalscope model is involved.

The primary evaluation was restricted to genes represented in the simulated Visium feature matrix and spots containing at least 50% malignant cells. For the set Ω of matched gene–spot pairs, reconstruction error was quantified using root mean squared error:

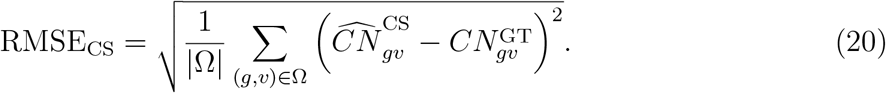

A constant-diploid predictor served as a baseline:

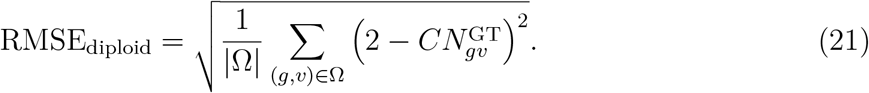

Both errors were calculated over identical gene–spot pairs. Relative improvement over the diploid baseline was defined as

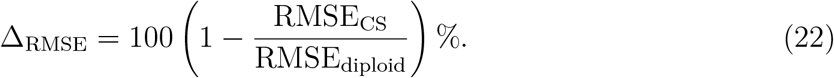

We additionally evaluated all spots and the subset of gene–spot pairs with substantial simulated copy-number deviation, 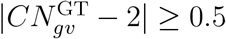. The latter analysis assesses reconstruction specifically in regions exhibiting appreciable copy-number alterations.

#### Copy-number signal concordance using CopyKAT

CopyKAT [11] was independently applied to the simulated spatial expression matrices. Its saved CNA matrix contains continuous, normalized copy-number-associated signals rather than directly calibrated absolute total-copy-number estimates.

Accordingly, CopyKAT was evaluated in its native signal representation without introducing an empirically fitted transformation to absolute copy number.

CopyKAT genomic positions were aligned with simulated gene coordinates using the nearest position on the same chromosome. Gene assignments were retained when the midpoint-to-position distance was at most 500 kb. Evaluation was restricted to genes represented in the simulated Visium feature matrix and spots with available CopyKAT CNA estimates.

Let *S*_*gv*_ denote the CopyKAT signal assigned to gene *g* in spot *v*, and define the ground-truth copy-number deviation from diploid as

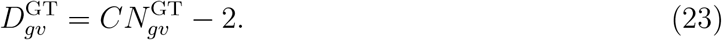

Continuous CNA-signal concordance was assessed using Pearson and Spearman correlations between the method-specific signal (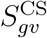 for Clonalscope or *S*_*gv*_ for CopyKAT) and 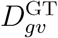 over matched gene–spot observations. Both correlations are calculated on the available matched observations for each method, without comparing the methods’ raw signal magnitudes.

We also evaluated discrimination between copy-number gains and losses. A ground-truth gain was defined by 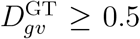, and a loss by 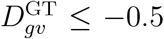. The area under the receiver operating characteristic curve (AUROC) was calculated using each method’s continuous deviation signal to distinguish gains from losses among these altered observations. Neutral observations with 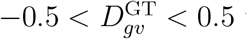 were excluded from this binary discrimination metric.

Directional agreement was defined as

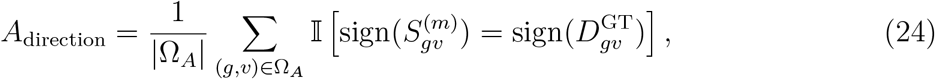

where Ω_*A*_ contains gene–spot pairs with ground-truth gains or losses, 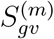 denotes either the Clonalscope centered signal 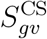 or the native CopyKAT signal *S*_*gv*_, and I[·] denotes the indicator function.

Gain-versus-loss AUROC and directional agreement therefore characterize CNA signal orientation and discrimination rather than calibrated absolute copy-number reconstruction. Clonalscope was additionally evaluated with absolute copy-number RMSE under the diploid calibration in Eq. (18); no absolute total-copy-number RMSE was assigned to CopyKAT’s uncalibrated CNA signal. For each method, copy-number profile availability was defined as the proportion of all simulated spatial spots for which a genomic CNA profile was returned, regardless of any binary malignant/normal classification.

#### Shared-observation evaluation

To account for differences in genomic coverage and available spot predictions between Clonalscope and CopyKAT, an additional comparison was performed on their intersecting gene and spot sets.

Genes were retained when they were represented in the simulated Visium expression matrix and could be assigned to genomic regions in both methods. Spots were retained when both methods returned CNA estimates.

Clonalscope reconstruction RMSE and CopyKAT CNA-signal concordance were previously recalculated on identical gene–spot observations. For the aligned signal comparison, both methods’ centered signals are evaluated using the same correlation, gain-versus-loss AUROC, and direction-agreement measures on their matching observations. This shared-observation evaluation distinguishes differences caused by the available genomic and spatial observations from differences in signal reconstruction.

The shared evaluation comprised 99 spots and 22,012 genes for HGSOC, 200 spots and 17,947 genes for CRC, and 211 spots and 15,992 genes for HCC.

Because Clonalscope and CopyKAT return different numerical representations, common signal-based metrics are interpreted as concordance with ground truth rather than direct calibration of the two numerical signal scales. Clonalscope’s absolute-copy-number RMSE remains a separate, diploid-calibrated analysis and is not equated with CopyKAT correlations.

#### Malignant spot classification and subclone recovery

Ground-truth malignant-spot status was defined by a malignant-cell fraction of at least 0.5. Binary predictions from downstream methods were compared with these labels using precision, recall, specificity, and F1 score. The percentage of spots receiving a defined binary malignant/normal classification was reported separately, and classification metrics were calculated only on spots with an explicit tumor or normal assignment. Undefined or missing labels were excluded rather than recoded into either class.

Clonalscope malignant subclone recovery was evaluated by comparing inferred cluster labels with the dominant ground-truth malignant subclone within each spot. The primary analysis included spots with malignant fraction at least 0.5 and dominant-subclone fraction at least 0.5 among malignant cells. A stricter dominant-subclone threshold of 0.8 was also evaluated.

Clustering agreement was summarized using the adjusted Rand index (ARI), V-measure, homogeneity, and completeness. These metrics are invariant to permutations of inferred cluster labels and do not require a predefined one-to-one correspondence between predicted clusters and true malignant subclones. For each evaluated dominance threshold, the number of distinct Clonalscope cluster IDs among included spots was also counted and compared with the number of distinct ground-truth malignant subclone labels. This count uses the unsupervised cluster labels rather than Clonalscope’s subsequent binary tumor/normal classification.

### Evaluation and comparison of malignant subclone spatial organization

#### Spatial autocorrelation of OncoSpot malignant subclones

Spatial autocorrelation was evaluated using the final cell-level spatial coordinates and malignant subclone assignments generated by OncoSpot. The analysis included 7,500 malignant cells from each of the HGSOC, CRC, and HCC datasets; nonmalignant cells were excluded. Malignant subclone identities corresponded to the final retained subclones after phylogeny-guided merging of rare lineages.

For each dataset, we constructed a symmetrized eight-nearest-neighbor graph using Euclidean distances between malignant-cell coordinates. Duplicate edges were removed, yielding an undirected spatial graph with binary adjacency weights *w*_*ij*_ ∈ {0, 1}.

Spatial clustering was quantified using same-clone join-count statistics and Moran’s *I* [25].

The same-clone join fraction was defined as

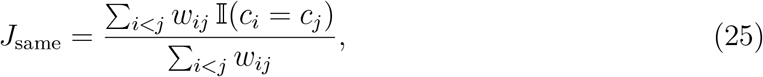

where *w*_*ij*_ indicates spatial adjacency between cells *i* and *j*, and *c*_*i*_ denotes the malignant subclone of cell *i*.

Under random reassignment of clone identities while preserving clone abundances, the expected same-clone join fraction is

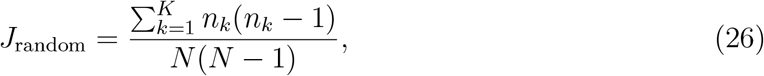

where *K* is the number of retained malignant subclones, *n*_*k*_ denotes the number of cells belonging to subclone *k*, and *N* is the total number of malignant cells. Join-count enrichment was calculated as

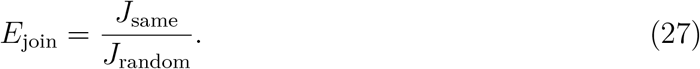

For each subclone *k*, Moran’s *I* was calculated using the binary indicator 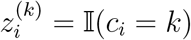:

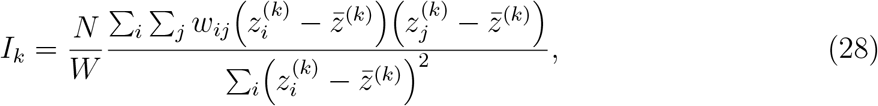

where *W* = Σ_*i*_ Σ_*j*_ *w*_*ij*_. The primary summary statistic was the median Moran’s *I* across retained subclones representing at least 1% of malignant cells.

Statistical significance was assessed using 999 random permutations of malignant subclone identities while preserving clone abundances, spatial coordinates, and graph connectivity. One-sided permutation tests assessed enrichment of same-clone joins and positive Moran’s *I*. For the clone-specific Moran’s *I* tests, *P* -values were adjusted using the Benjamini– Hochberg procedure. Permutation *P* -values were calculated with a plus-one correction, giving a minimum attainable value of 0.001.

#### Phylogeny–space concordance of malignant subclones

To evaluate whether spatially adjacent malignant subclones are also closely related on the evolutionary tree, we generated 50 independent OncoSpot realizations for each of HGSOC, CRC, and HCC, using distinct random seeds and the corresponding cancer-specific simulation configurations. Each realization comprised 15,000 cells, including 7,500 malignant cells, and an independently generated malignant subclone phylogeny and Visium-like spatial observation. Malignant subclone identities corresponded to the final retained clones after phylogeny-guided merging of rare lineages.

Phylogeny–space concordance was evaluated at the Visium spot level. For each spot containing malignant cells, the dominant malignant subclone was assigned only if it represented at least 80% of the malignant cells captured in that spot. Spots without malignant cells or without a sufficiently dominant malignant subclone were excluded from boundary evaluation.

No additional threshold on the malignant fraction of all captured cells was imposed. Spatial adjacency was defined by direct nearest-neighbor connections on the hexagonally staggered Visium spot grid. The adjacency graph was constructed using all occupied spots before restricting edges to pairs of eligible spots assigned to different malignant subclones.

Let *ℰ* denote the set of qualifying cross-subclone edges, and let *c*(*v*) denote the dominant malignant subclone assigned to spot *v*. We defined the phylogeny–space boundary statistic as

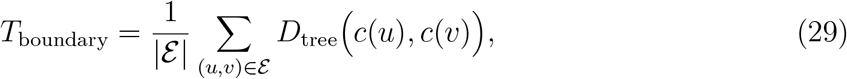

where *D*_tree_(*a, b*) is the unweighted shortest-path distance, measured in numbers of edges, between subclone nodes *a* and *b* on the retained malignant phylogeny. Ancestral nodes were included when calculating tree distances. Smaller values of *T*_boundary_ indicate that spatially adjacent subclones tend to be more closely related evolutionarily.

For each realization, we constructed a null distribution by permuting the identities of the final observed malignant subclones among their corresponding nodes on the fixed evolutionary tree. This procedure preserved the spatial arrangement of subclones, spot-level clone assignments, spatial adjacency, clone abundances, and tree topology, while disrupting the correspondence between spatial clone identities and phylogenetic positions. Ancestor-only nodes without retained malignant-cell populations remained fixed as structural nodes. All possible permutations were enumerated when their number did not exceed 1,000; otherwise, 1,000 distinct random permutations were sampled using a reproducible random seed.

The concordance score for each realization was standardized against its permutation null distribution:

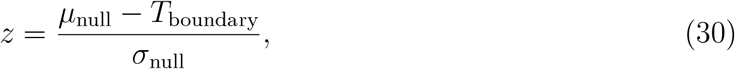

where *µ*_null_ and *σ*_null_ are the mean and population standard deviation of the null boundary statistics, respectively. Positive *z*-scores indicate that observed neighboring subclones are phylogenetically closer than expected under randomized clone–tree correspondence.

Realizations lacking the required simulation outputs, containing fewer than two retained malignant subclones, having no qualifying cross-subclone boundaries, or producing zero null standard deviation were excluded from the standardized analysis because a valid *z*-score could not be calculated. All other realizations were retained regardless of the direction of their *z*-scores. For each cancer, the resulting *z*-scores were summarized by their median and interquartile range. A one-sided one-sample Wilcoxon signed-rank test was used to assess whether the distribution of *z*-scores was shifted above zero.

#### Reference-free spatial-pattern comparison with scCube

To compare malignant subclone spatial organization with an established spatial transcriptomics simulator, we evaluated the reference-free spatial-pattern generator of scCube [19]. The comparison was designed to isolate spatial placement under matched malignant clone populations rather than compare the complete transcriptomic simulation pipelines.

For each cancer, the 7,500 malignant cells generated by OncoSpot were used to define the cell population supplied to scCube. Final post-merge malignant subclone identities were treated as population labels, with exact clone abundances preserved. The resulting populations comprised five subclones for HGSOC, six for CRC, and five for HCC.

The scCube `generate_pattern_random()` function was used to produce two-dimensional reference-free spatial coordinates without modifying the original scCube implementation. The default spatial-pattern parameters were retained: spatial size 30, distance-covariance parameter *δ* = 25, and coordinate-swapping parameter *λ* = 0.75. Spatial dimensionality was set to two. The spatial-pattern function was supplied with the matched cell identifiers and clone annotations; VAE training and expression generation were omitted because the evaluation specifically targeted spatial organization rather than transcriptomic fidelity.

For each cancer, ten independent scCube spatial realizations were generated using predefined random seeds 20261027 through 20261036. Each realization preserved the exact identities and abundances of the corresponding malignant clone populations. Generated spatial coordinates were validated for completeness, uniqueness, and finite numerical values, and clone assignments were checked against the corresponding OncoSpot annotations.

The same spatial autocorrelation evaluation was then applied independently to each scCube realization. Specifically, a symmetrized eight-nearest-neighbor graph was constructed from the generated coordinates, and same-clone join enrichment and per-clone Moran’s *I* were calculated using Eqs. (25)–(28). Statistical significance was assessed using 999 clone-label permutations per realization, with Benjamini–Hochberg adjustment for clone-specific tests. Identical statistical function implementations and neighborhood parameters were used for both simulators.

For scCube, spatial autocorrelation metrics were summarized by the median and interquartile range across the ten independent realizations. OncoSpot was represented by one finalized spatial realization per cancer; therefore, between-realization variability was characterized for scCube only. For visualization, the scCube realization with join enrichment closest to the median of its ten realizations was selected for each cancer.

The comparison preserves malignant clone abundances and evaluation definitions but does not impose identical tissue geometries or spatial-boundary constraints. OncoSpot coordinates reflect its deme-based evolutionary scaffold and subsequent spatial refinement, whereas scCube generates a reference-free spatial coordinate field. Accordingly, differences in spatial autocorrelation quantify differences between spatial-pattern generation mechanisms under matched clone compositions, rather than directly measure biological realism or independent transcriptomic simulation accuracy.

#### Phylogeny–space concordance comparison with scCube

To compare the relationship between evolutionary ancestry and spatial organization across simulators, scCube reference-free spatial coordinates were generated for the malignant-cell populations associated with the 124 OncoSpot realizations having evaluable native-Visium phylogeny–space scores. For each matched realization, scCube was supplied with the same 7,500 malignant-cell identities, final subclone labels, and clone abundances, without the OncoSpot phylogeny or clone-associated expression information. The corresponding OncoSpot tree was used only during evaluation. To make the spatial comparison consistent despite different native coordinate systems, we constructed hexagonally staggered pseudo-spot grids independently from the malignant-cell coordinates of both simulators. Grid spacing was calibrated without using clone labels, targeting approximately ten malignant cells per occupied pseudo-spot, and the capture-radius-to-spacing ratio was fixed at 0.275. Eligible pseudo-spots required a dominant malignant subclone representing at least 80% of their captured cells. Cross-subclone boundaries between directly adjacent eligible pseudo-spots were scored using Eq. (29) and the same clone-identity permutation null model (at most 1,000 permutations) used for OncoSpot. The primary comparison used one predefined grid origin per matched tree and a one-sided paired Wilcoxon signed-rank test of the difference *z*_OncoSpot_ −*z*_scCube_. Four additional grid offsets were examined as sensitivity analyses, not treated as independent realizations. Pairs without a defined score for both methods were excluded from paired tests. This malignant-cell-only pseudo-spot comparison was conducted separately from the native-Visium concordance analysis, which also includes nonmalignant cells in its spot construction.

### Datasets and cancer-specific parameterization

#### Reference datasets

OncoSpot is parameterized for HGSOC, CRC, and HCC using public single-cell RNA-seq, matched cancer genomic/transcriptomic resources, and spatial transcriptomic data (**Table 9**). Reference datasets play different roles for the malignant cell of origin, malignant transcriptional states and CNA response, nonmalignant cell types, and the Visium observation model. These components are independently calibrated where their source datasets differ, while the mathematical model and six state definitions remain common across cancers.

**Table 9:** Reference datasets and molecular resources for HGSOC, CRC, and HCC parameterization. Source-cell subsets and selection criteria are specified in the text and Supplemental Section S1.

| Cancer | Accession/resource | Data type | Parameterization role |
|---|---|---|---|
| HGSOC | GSE151214 (Dinh et al.) [41, 42] | Single-cell RNA-seq | Early-secretory fallopian-tube epithelial cell-of-origin baseline. |
| HGSOC | SA1053 (Funnell et al.; TreeAlign) [43, 9] | Matched tumor DNA/RNA | Malignant state-vector reference, gene-level state effects, malignant library sizes, and retained CNA-response/dispersion parameters. |
| HGSOC | GSE184880 (Xu et al.) [44, 45] | Single-cell RNA-seq | Nonmalignant expression and tumor-microenvironment reference populations. |
| Shared | GSE211956 SP4 (Denisenko et al.) [4, 40] | Visium | Empirical spot-level UMI-depth histogram for the observation model. |
| CRC | GSE132465 (Lee et al.) [46, 47] | Single-cell RNA-seq | Normal colonic epithelial origin, primary-tumor epithelial state and malignant-expression fits, and nonmalignant cell-type references. |
| HCC | GSE149614 (Lu et al.) [48, 49] | Single-cell RNA-seq | Normal-site hepatocyte origin, tumor-site hepatocyte state and malignant-expression fits, and nonmalignant cell-type references. |
| Shared | GENCODE v50 [50]; MSigDB Hallmark [39] | Gene annotation; gene sets | CNA-to-gene mapping and the six shared malignant transcriptional axes. |

#### HGSOC reference and evolutionary configuration

An early-secretory fallopian-tube epithelial population from GSE151214 supplies the HGSOC cell-of-origin baseline [41, 42]. The matched single-cell HGSOC resources associated with the SA1053 patient-derived xenograft and its TreeAlign assignments provide the malignant tumor reference [43, 9]. Exactly 3,369 expression-profiled cells with matched TreeAlign assignment, representing three samples from one biological patient, are used for six-dimensional Hallmark score estimation and joint-state resampling. The same reference counts support centered ridge estimation of gene-specific state-response coefficients. CNA-response and NB dispersion parameters are retained from the separately calibrated HGSOC expression parameterization. Nonmalignant expression is informed by GSE184880 [44, 45]. The shared Visium UMI-depth model is derived from sample SP4 of GSE211956 [4, 40].

The invasive-glandular DEMON preset represents a spatial invasion scenario motivated by HGSOC histology and documented regional clonal divergence [33–35]. Model selection does not imply that the DEMON dispersal mechanism describes every HGSOC tumor. Growth parameters were calibrated by repeated stochastic realizations without imposing a fixed final clone number. Calibration coverage, signature overlap, and retained gene-level parameters are summarized in the **Supplemental Sections S4-S5**.

#### CRC reference and evolutionary configuration

CRC parameters use GSE132465, which profiles primary colorectal tumors and matched normal mucosa [46, 47]. Malignant expression programs and their interactions with the tumor microenvironment have also been characterized in independent single-cell and spatial studies of CRC [51]. The normal epithelial reference was used to estimate a donor-balanced colon stem-like/transit-amplifying origin composition, while selected primary-tumor epithelial cells were used to estimate tumor-state distributions, empirical malignant library sizes, gene-level state effects, and dispersion. Gland fission is motivated by direct crypt-fission observations in colorectal adenomas, with applicability to CRC interpreted as a growth scenario rather than a universal mechanism [36, 23]. The DEMON preset values are documented in **Supplemental Section S2**. Nonmalignant expression parameters are estimated by broad reference cell type and donor with the same NB parameterization used for HCC. Spatial-compartment proportions and the shared observation settings are simulation inputs rather than direct estimates of CRC-specific tissue purity or capture efficiency.

#### HCC reference and evolutionary configuration

HCC expression parameters use GSE149614, a single-cell atlas containing primary tumors, adjacent non-tumor liver, portalvein tumor thrombi, and metastatic lymph-node specimens [48, 49]. Normal-site hepatocytes were selected for the donor-balanced cell-of-origin baseline. Tumor-site cells labeled “Hepatocyte” supply the empirical malignant-reference library distribution and tumor-state/expression calibration. Site and broad cell-type labels were used as the selection criteria: independent malignant-genotype confirmation of every selected tumor-site hepatocyte was not available, and the resulting calibration is therefore provisional. Primary-tumor nonmalignant populations inform six broad normal classes, with normal-site hepatocytes providing the normal-epithelial class.

Gene symbols were mapped to autosomal GENCODE v50 gene identifiers by unique matches [50]. Uniquely mapped autosomal genes were retained. The retained gene universe and full CNA-coverage audit for HCC are detailed in the **Supplemental Sections S4-S5**. The six Hallmark axes are scored in tumor-site hepatocytes, giving HCC-specific empirical joint-state vectors and gene-level response coefficients. The boundary-restricted DEMON preset represents selected expansile HCC growth patterns rather than all HCC morphologies [37, 38, 23]. Its numerical settings are given in **Supplemental Section S2**. CNAsim, spatial placement, and Visium observation are independently specified simulation modules, not measurements of HCC-specific tissue growth or capture efficiency. In addition, a subset of normal-site hepatocyte reference cells exhibited unexpectedly high immunoglobulin counts. These were documented as a reference-quality caveat rather than silently removed from the present baseline.

#### Reference-processing and parameter status

Across HGSOC, CRC, and HCC, the six transcriptional axes and their scoring procedure are identical, but empirical state distributions, fitted state effects, cell-of-origin baselines, library-size distributions, and expression dispersion are cancer-dependent. HGSOC state coefficients are globally centered because its malignant reference contains one biological patient, whereas CRC and HCC use within-patient centering. HGSOC also retains gene-dependent CNA dosage parameters, whereas CRC and HCC use a fixed dosage exponent. Exact reference subsets, parameter values, and gene coverage are reported in **Supplemental Sections S2, S4-S5**.

## Data and code availability

OncoSpot is freely available at https://github.com/compbio-mallory/OncoSpot.

## Acknowledgment

This work was supported by NSF CCF grant number 2523717 to Xian Mallory, the NIH NIGMS Maximizing Investigators’ Research Award (MIRA) R35 GM146960 to Xin Maizie Zhou, and Waddell Walker Hancock Cancer Discovery Fund Award to Xin Maizie Zhou.

## References

[1] Patrik L. Ståhl, Fredrik Salmén, Sanja Vickovic, Anna Lundmark, José Fernández Navarro, Jens Magnusson, Stefania Giacomello, Michaela Asp, Jakub O. Westholm, Mikael Huss, Annelie Mollbrink, Sten Linnarsson, Simone Codeluppi, Åke Borg, Fredrik Pontén, Paul Igor Costea, Pelin Sahlén, Jan Mulder, Olaf Bergmann, Joakim Lundeberg, and Jonas Frisén. Visualization and analysis of gene expression in tissue sections by spatial transcriptomics. Science, 353(6294):78–82, 2016. doi: 10.1126/science.aaf2403.

[2] Artem Lomakin, Jessica Svedlund, Carina Strell, et al. Spatial genomics maps the structure, nature and evolution of cancer clones. Nature, 611:594–602, 2022. doi: 10.1038/s41586-022-05425-2.

[3] Tongtong Zhao, Zachary D. Chiang, Julia W. Morriss, et al. Spatial genomics enables multi-modal study of clonal heterogeneity in tissues. Nature, 601:85–91, 2022. doi: 10.1038/s41586-021-04217-4.

[4] Elena Denisenko, Leanne de Kock, Adeline Tan, et al. Spatial transcriptomics reveals discrete tumour microenvironments and autocrine loops within ovarian cancer subclones. Nature Communications, 15:2860, 2024. doi: 10.1038/s41467-024-47271-y.

[5] Rebecca Elyanow, Ron Zeira, Max Land, and Benjamin J. Raphael. STARCH: copy number and clone inference from spatial transcriptomics data. Physical Biology, 18(3): 035001, 2021. doi: 10.1088/1478-3975/abbe99.

[6] Cong Ma, Metin Balaban, Jingxian Liu, Siqi Chen, Michael J. Wilson, Christopher H. Sun, Li Ding, and Benjamin J. Raphael. Inferring allele-specific copy number aberrations and tumor phylogeography from spatially resolved transcriptomics. Nature Methods, 21 (12):2239–2247, 2024. doi: 10.1038/s41592-024-02438-9.

[7] Dalia Barkley, Reuben Moncada, Maayan Pour, et al. Cancer cell states recur across tumor types and form specific interactions with the tumor microenvironment. Nature Genetics, 54:1192–1201, 2022. doi: 10.1038/s41588-022-01141-9.

[8] Avishai Gavish, Michael Tyler, Alissa C Greenwald, Rouven Hoefflin, Dor Simkin, Roi Tschernichovsky, Noam Galili Darnell, Einav Somech, Chaya Barbolin, Tomer Antman, et al. Hallmarks of transcriptional intratumour heterogeneity across a thousand tumours. Nature, 618(7965):598–606, 2023.

[9] Hongyu Shi, Marc J. Williams, Gryte Satas, Adam C. Weiner, Andrew McPherson, and Sohrab P. Shah. Allele-specific transcriptional effects of subclonal copy number alterations enable genotype-phenotype mapping in cancer cells. Nature Communications, 15:2482, 2024. doi: 10.1038/s41467-024-46710-0.

[10] Chi-Yun Wu, Jiazhen Rong, Anuja Sathe, Paul R. Hess, Billy T. Lau, Susan M. Grimes, Sijia Huang, Hanlee P. Ji, and Nancy R. Zhang. Cancer subclone detection based on DNA copy number in single-cell and spatial omic sequencing data. Nature Methods, 22 (9):1846–1856, 2025. doi: 10.1038/s41592-025-02773-5.

[11] Ruli Gao, Shanshan Bai, Ying C. Henderson, Yiyun Lin, Aislyn Schalck, et al. Delineating copy number and clonal substructure in human tumors from single-cell transcriptomes. Nature Biotechnology, 39(5):599–608, 2021. doi: 10.1038/s41587-020-00795-2.

[12] Zhixing Zhong, Junchen Hou, Zhixian Yao, Lei Dong, Feng Liu, et al. Domain generalization enables general cancer cell annotation in single-cell and spatial transcriptomics. Nature Communications, 15:1929, 2024. doi: 10.1038/s41467-024-46413-6.

[13] Qiaoni Yu, Yuan-Yuan Li, and Yunqin Chen. scMalignantFinder distinguishes malignant cells in single-cell and spatial transcriptomics by leveraging cancer signatures. Communications Biology, 8:504, 2025. doi: 10.1038/s42003-025-07942-y.

[14] Pankaj Bhattarai, Weiman Yuan, Hongmei Chi, Xin Maizie Zhou, and Xian Mallory. Multi-modality graph representation learning for malignant cell identification from scRNA-seq using DeepMalignant. bioRxiv, 2026. doi: 10.64898/2026.06.29.734828. Preprint.

[15] Dylan M. Cable, Evan Murray, Luli S. Zou, Aleksandrina Goeva, Evan Z. Macosko, Fei Chen, and Rafael A. Irizarry. Robust decomposition of cell type mixtures in spatial transcriptomics. Nature Biotechnology, 40(4):517–526, 2022. doi: 10.1038/s41587-021-00830-w.

[16] Vitalii Kleshchevnikov, Artem Shmatko, Emma Dann, Alexander Aivazidis, Hamish W. King, et al. Cell2location maps fine-grained cell types in spatial transcriptomics. Nature Biotechnology, 40:661–671, 2022. doi: 10.1038/s41587-021-01139-4.

[17] Shi-Min Han, Zhi-Xing Xiong, Ying-Shi Zhou, and Can Yang. Benchmarking copy number alteration inference methods for spatial transcriptomics. Nature Communications, 17(1): 10565, 2026. doi: 10.1038/s41467-026-77500-5.

[18] Jiaqiang Zhu, Lu-Lu Shang, and Xiang Zhou. SRTsim: spatial pattern preserving simulations for spatially resolved transcriptomics. Genome Biology, 24(1):39, 2023. doi: 10.1186/s13059-023-02879-z.

[19] Jingyang Qian, Hudong Bao, Xin Shao, Yin Fang, Jie Liao, Zhuo Chen, et al. Simulating multiple variability in spatially resolved transcriptomics with scCube. Nature Communications, 15:5021, 2024. doi: 10.1038/s41467-024-49445-0.

[20] Hongrui Duo, Yinghong Li, Yang Lan, Jingxin Tao, Qingxia Yang, et al. Systematic evaluation with practical guidelines for single-cell and spatially resolved transcriptomics data simulation under multiple scenarios. Genome Biology, 25:145, 2024. doi: 10.1186/s13059-024-03290-y.

[21] Shen-Jie Wang, Yu-Hang Li, Xiao-Nan Wang, Xu-Wen Wang, Tian-Ci Wang, Shuan-Ying Yang, and Jia-Yin Wang. STGBench: sequencing-level spatial DNA–RNA simulation for multimodal and virtual cell-oriented benchmarking of genomic alterations. Briefings in Bioinformatics, 27:bbag354, 2026. doi: 10.1093/bib/bbag354.

[22] Xianjie Huang, Rongting Huang, Jiamu Qiao, and Yuan-Hua Huang. stCNASim: allele-aware spatial RNA-seq simulator enables systematic benchmarking of copy number inference, 2026.

[23] Robert Noble, Dominik Burri, Cécile Le Sueur, Jeanne Lemant, Yannick Viossat, Jakob Nikolas Kather, and Niko Beerenwinkel. Spatial structure governs the mode of tumour evolution. Nature Ecology & Evolution, 6:207–217, 2022. doi: 10.1038/s41559-021-01615-9.

[24] Samson Weiner and Mukul S. Bansal. CNAsim: improved simulation of single-cell copy number profiles and DNA-seq data from tumors. Bioinformatics, 39(7):btad434, 2023. doi: 10.1093/bioinformatics/btad434.

[25] Patrick A. P. Moran. Notes on continuous stochastic phenomena. Biometrika, 37(1–2): 17–23, 1950. doi: 10.1093/biomet/37.1-2.17.

[26] Cancer Genome Atlas Network1 et al. Comprehensive molecular characterization of human colon and rectal cancer. Nature, 487(7407):330, 2012.

[27] Adrian Ally, Miruna Balasundaram, Rebecca Carlsen, Eric Chuah, Amanda Clarke, Noreen Dhalla, Robert A Holt, Steven JM Jones, Darlene Lee, Yussanne Ma, et al. Comprehensive and integrative genomic characterization of hepatocellular carcinoma. Cell, 169(7):1327–1341, 2017.

[28] Pankaj Bhattarrai, Weiman Yuan, Hongmei Chi, Xin Maizie Zhou, and Xian Mallory. Multi-modality graph representation learning for malignant cell identification from scrna-seq using deepmalignant. bioRxiv, pages 2026–06, 2026.

[29] Cancer Genome Atlas Research Network et al. Integrated genomic analyses of ovarian carcinoma. Nature, 474(7353):609, 2011.

[30] Rosalin A. Cooper, Emily Thomas, Anna M. Sozanska, Carlo Pescia, and Daniel J. Royston. Spatial transcriptomic approaches for characterising the bone marrow landscape: pitfalls and potential. Leukemia, 39:291–295, 2025. doi: 10.1038/s41375-024-02480-8.

[31] Christopher P. Ly, Ivo Veletic, Christopher D. Pacheco, et al. Multimodal spatial proteomic profiling in acute myeloid leukemia. npj Precision Oncology, 9:148, 2025. doi: 10.1038/s41698-025-00897-7.

[32] Amanda Janesick, Robert Shelansky, Andrew D. Gottscho, Florian Wagner, Stephen R. Williams, Morgane Rouault, Ghezal Beliakoff, et al. High resolution mapping of the tumor microenvironment using integrated single-cell, spatial and in situ analysis. Nature Communications, 14:8353, 2023. doi: 10.1038/s41467-023-43458-x.

[33] Russell Vang, Ie-Ming Shih, and Robert J. Kurman. Ovarian low-grade and highgrade serous carcinoma: pathogenesis, clinicopathologic and molecular biologic features, and diagnostic problems. Advances in Anatomic Pathology, 16(5):267–282, 2009. doi: 10.1097/PAP.0b013e3181b4fffa.

[34] Amy B. Bromley, Alon D. Altman, Pamela Chu, Jill G. Nation, Gregg S. Nelson, Praful Ghatage, Steve E. Kalloger, Guangming Han, and Martin Köbel. Architectural patterns of ovarian/pelvic high-grade serous carcinoma. International Journal of Gynecological Pathology, 31(5):397–404, 2012. doi: 10.1097/PGP.0b013e31824c2372.

[35] Andrew McPherson, Andrew Roth, Ewa Laks, et al. Divergent modes of clonal spread and intraperitoneal mixing in high-grade serous ovarian cancer. Nature Genetics, 48: 758–767, 2016. doi: 10.1038/ng.3573.

[36] W. M. Wong, N. Mandir, R. A. Goodlad, B. C. Y. Wong, S. B. Garcia, S. K. Lam, and N. A. Wright. Histogenesis of human colorectal adenomas and hyperplastic polyps: the role of cell proliferation and crypt fission. Gut, 50(2):212–217, 2002. doi: 10.1136/gut.50.2.212.

[37] T. Nakashima, M. Kojiro, Y. Kawano, F. Shirai, N. Takemoto, Y. Tomimatsu, H. Kawasaki, and K. Okuda. Histologic growth pattern of hepatocellular carcinoma: relationship to orcein (hepatitis B surface antigen)-positive cells in cancer tissue. Human Pathology, 13(6):563–568, 1982. doi: 10.1016/S0046-8177(82)80272-4.

[38] Manuel Schlageter, Luigi Maria Terracciano, Salvatore D’Angelo, and Paolo Sorrentino. Histopathology of hepatocellular carcinoma. World Journal of Gastroenterology, 20(43): 15955–15964, 2014. doi: 10.3748/wjg.v20.i43.15955.

[39] Arthur Liberzon, Chet Birger, Helga Thorvaldsdóttir, Mahmoud Ghandi, Jill P. Mesirov, and Pablo Tamayo. The molecular signatures database (MSigDB) hallmark gene set collection. Cell Systems, 1(6):417–425, 2015. doi: 10.1016/j.cels.2015.12.004.

[40] NCBI Gene Expression Omnibus. GSE211956: Spatial transcriptomics data associated with ovarian cancer subclone microenvironments. https://www.ncbi.nlm.nih.gov/geo/query/acc.cgi?acc=GSE211956, 2022. GEO Series accession GSE211956.

[41] Huy Q. Dinh, Xianzhi Lin, Farnaz Abbasi, et al. Single-cell transcriptomics identifies gene expression networks driving differentiation and tumorigenesis in the human fallopian tube. Cell Reports, 35(2):108978, 2021. doi: 10.1016/j.celrep.2021.108978.

[42] NCBI Gene Expression Omnibus. GSE151214: Single-cell transcriptomics identifies transcriptional networks driving differentiation and tumorigenesis in the human fallopian tube. https://www.ncbi.nlm.nih.gov/geo/query/acc.cgi?acc=GSE151214, 2021. GEO Series accession GSE151214.

[43] Tyler Funnell, Ciara H. O’Flanagan, Marc J. Williams, et al. Single-cell genomic variation induced by mutational processes in cancer. Nature, 612:106–115, 2022. doi: 10.1038/s41586-022-05249-0.

[44] Junfen Xu, Yifeng Fang, Kelie Chen, Sen Li, Sangsang Tang, et al. Single-cell RNA sequencing reveals the tissue architecture in human high-grade serous ovarian cancer. Clinical Cancer Research, 28(16):3590–3602, 2022. doi: 10.1158/1078-0432.CCR-22-0296.

[45] NCBI Gene Expression Omnibus. GSE184880: Single-cell rna sequencing reveals tissue architecture during human high-grade serous ovarian cancer progression. https://www.ncbi.nlm.nih.gov/geo/query/acc.cgi?acc=GSE184880, 2022. GEO Series accession GSE184880.

[46] Hae-Ock Lee, Yourae Hong, Hakki Emre Etlioglu, et al. Lineage-dependent gene expression programs influence the immune landscape of colorectal cancer. Nature Genetics, 2020. doi: 10.1038/s41588-020-0636-z.

[47] NCBI Gene Expression Omnibus. GSE132465: Single cell 3’ rna sequencing of 23 korean colorectal cancer patients. https://www.ncbi.nlm.nih.gov/geo/query/acc.cgi?acc=GSE132465, 2020. GEO Series accession GSE132465.

[48] Yiming Lu, Aiqing Yang, Cheng Quan, Yingwei Pan, et al. A single-cell atlas of the multicellular ecosystem of primary and metastatic hepatocellular carcinoma. Nature Communications, 13:4594, 2022. doi: 10.1038/s41467-022-32283-3.

[49] NCBI Gene Expression Omnibus. GSE149614: A single-cell atlas of the multicellular ecosystem of primary and metastatic hepatocellular carcinoma. https://www.ncbi.nlm.nih.gov/geo/query/acc.cgi?acc=GSE149614, 2020. GEO Series accession GSE149614.

[50] GENCODE Consortium. GENCODE human release 50 (GRCh38.p14). https://www.gencodegenes.org/human/, 2026. Human gene annotation release 50.

[51] Teng Wang, Zhaoming Chen, Wang Wang, Heng Wang, and Shenglong Li. Single-cell and spatial transcriptomic analysis reveals tumor cell heterogeneity and underlying molecular program in colorectal cancer. Frontiers in Immunology, 16:1556386, 2025. doi: 10.3389/fimmu.2025.1556386.

